# Leaving X: How scale insects evolved alternatives to chromosomal sex determination

**DOI:** 10.64898/2026.01.31.703042

**Authors:** Andrew J. Mongue, Erin C. Powell, Tracy Liesenfelt, Ethan Grebler, Amanda Markee

## Abstract

Reproduction is a core feature of life, which makes understanding the reproductive diversity of organisms a fundamental goal of evolutionary biology. It is still unclear why sex determination systems are faithfully conserved for hundreds of millions of years in some taxa but repeatedly turn over in others. A prime example is the true bugs, Hemiptera. The vast majority of families employ a conserved X sex chromosomal system, but in scale insects (Coccomorpha), sex determination systems are far more varied. Different families may retain X chromosomes, employ a rare form of haplodiploidy known as paternal genome elimination (PGE), or forgo sexual reproduction in favor of simultaneous hermaphroditism. While these systems have been known for decades, only recently has non-model genomics enabled their study with modern bioinformatic methods. We collected and sequenced representatives of rare, early-diverging scale insects to study genomic changes that coincide with transitions between sex determination systems. Using ancillary expression data, we also explored sex-biased gene expression in a subset of scale insects on both sides of one transition. In a newly updated molecular phylogeny, we recovered two independent losses of X chromosomal sex determination, once in the family Monophlebidae and a second time at the base of the families that employ PGE. In the former, genomic rearrangements begin before the X is lost and any previously X-linked orthologs have been absorbed into one of two massive, shuffled autosomes in the hermaphrodites. In the latter, the X chromosome itself is faithfully preserved in early diverging PGE taxa; it appears to have merely lost its sex determination function. We analyze ortholog conservation across these transitions and find few genes lost, with only one orthogroup lost in both transitions, and a number of orthogroups gained, suggesting these large transitions did not require large changes to the gene content of the X chromosome. Expression data suggests that extreme sexual dimorphism of gene expression predates sex determination turnover and may have helped enable it. Together these new observations illustrate the multiple routes to sex determination turnover, help refine hypotheses for its causes, and provide a wealth of new resources for the modern genomic study of scale insects.

## Introduction

The core goal of evolutionary biology is to explain the patterns of diversity of life and the processes that generate these patterns. Across more than 150 years of research, traits involved in reproduction have been of particular interest (starting with Darwin 1871), but the true extent of their diversity was not fully apparent until scientists began to study microscopic traits, including the chromosomes themselves. The discovery of chromosomal sex determination grew out of observations of what were initially known as “accessory chromosomes,” so termed because they varied between males and females of the same species of insects and were thus initially inferred to not be strictly necessary for the organism (McClung 1901, 1902). In subsequent research, their true importance in sex determination has been recognized, as have the numerous variations on these systems (Bachtrog et al. 2014), including some that defy Mendel’s familiar laws of inheritance (Ross et al. 2022). This staggering array of reproductive diversity belies something of a paradox, however. Despite undeniable overall diversity, many ancient, speciose clades have sex determination systems that are faithfully conserved across hundreds of thousands of species and hundreds of millions of years of evolution (Rodríguez Delgado et al. 2009; Fraïsse et al. 2017; Vicoso 2019). Thus, a more specific goal of evolutionary reproductive biology is to understand how and under what conditions sex determination systems turn over.

To reach this goal requires systematic knowledge of sex determination systems across species, a way to track the genomic and genetic changes that enable it, and a suitable system in which to study the variation. Tools to assess the former have been available for decades thanks to microscopy and cytogenetic analyses. Note that the observations of sex chromosomes referenced above are more than a century old (McClung 1901) and subsequent studies have surveyed such traits broadly in the intervening decades (Hughes-Schrader 1925; Núñez 1962; White 1973). It is the second piece, a deeper understanding of the genomic and genetic changes that underlie differences in karyotype, that has lagged behind. The field of genome sequencing is barely more than 30 years old (Olson 1993) and extension of these approaches beyond model species is roughly half that age (Ekblom and Galindo 2011; Unamba et al. 2015), so it is only recently that it has become tractable to ask broadly how turnover of sex determination occurs at the genomic and genetic levels.

Finally, with both types of evidence cataloged or easily attainable, it is worth carefully considering the choice of study taxa. As already alluded to, sex determination is variable across the tree of life but tends to be conserved across large taxonomic groups. For instance, mammals generally share the same orthologous X sex chromosome across over 100 million years of evolution (Rodríguez Delgado et al. 2009) and, thus, offer limited insight into sex determination turnover. And even where sex chromosome systems have turned over, like-for-like transitions are the most commonly studied (Vicoso and Bachtrog 2015), for example the X chromosome being replaced by a new X chromosome that was formerly an autosome (Vicoso and Bachtrog 2013). While there is still more to learn in these systems, researchers have already done much to characterize these turnovers. For instance, theory predicts that sexual conflict can favor the evolution of new sex determination genes on autosomes, which can supplant the existing sex determination mechanism (Van Doorn and Kirkpatrick 2007). Likewise genomic analysis of like-for-like transitions in *Drosophila* flies has shown that such turnovers are associated with the movement of genes expressed in males off of new X sex chromosomes (Layana et al. 2025), as predicted based on the fact that X chromosomes spend more time in females than males (Klein et al. 2021). Results from these systems do little to explain other, non-like-for-like turnovers of sex determination, however.

### Scale insects and the diversity of reproductive strategies

Scale insects (Hemiptera: Coccomorpha) are a uniquely well-suited choice for the study of drastic sex determination turnover (Gavrilov 2007). This diverse infraorder, consisting of approximately 32 extant and 19 extinct families (Vea and Grimaldi 2016; Powell et al. 2024) of plant-feeding insects, belongs to the order Hemiptera (true bugs), which generally have well-conserved XX/X0 or XY sex determination (Pal and Vicoso 2015); in other words, most Hemiptera share X chromosomal sex determination and the same orthologous chromosome serves as the X across species. However, within scale insects, different families may retain chromosomal XX/X0 sex determination (Nur 1980), employ a rare form of haplodiploidy known as paternal genome elimination (Nur 1966; de la Filia et al. 2021), or even largely forgo sexual reproduction, as in the simultaneous hermaphrodite giant scale insects (Hughes-Schrader 1925), at least some of which still produce rare haploid males (Royer 1975; Hodgson and Foldi 2006; Mongue et al. 2021). In both cases, the X chromosome system has been replaced with a fully autosomal genome in which the dynamics of the female/hermaphrodite genome (*i.e.*, full diploidy) did not change across these transitions. Males, in contrast, have transitioned from mostly diploidy with a single haploid (X) chromosome, to genome-wide haploidy, albeit in two very different contexts. Under paternal genome elimination, offspring sex ratio may fluctuate over a mother’s lifespan but overall remains at the classical 50-50 male-to-female ratio (Ross et al. 2012a), but in the hermaphrodites, male production is rare, with the vast majority of individuals being self-fertile hermaphrodites that produce more self-fertile offspring (Mongue et al. 2021). Uncovering when and how scale insects have transitioned between these systems can shed crucial light on the causes and consequences of sex determination turnover occurs more generally.

To do so, we have (1) sequenced 17 previously unsampled species and assembled genomes for each, including chromosome-level genomes for five species of scale insects from different families and sex determination systems, (2) confirmed their places in the evolution of this group via phylogenomics, (3) performed macrosynteny analyses to understand patterns of conservation or change of chromosome organization, examined (4) orthology across scales to track conservation of X-linked genes, and (5) expression differences between the sexes of some species for insight into the state of the X chromosome before and after transitions.

Together these results show that the otherwise conserved hemipteran X has been lost twice, in one case absorbed into a much larger autosome in the hermaphroditic scales as part of a massive condensing and shuffling of chromosomes and, in the other case of paternal genome elimination, the X chromosome was retained essentially intact at first, but apparently without its sex determining function. Thus, major turnovers can occur in distinct ways, even from the same starting genomic architecture.

## Methods

### Sourcing representatives of different sex determination systems

We downloaded existing datasets for scale insects representing different sex determination and reproductive systems. These data ranged from assembled chromosome-level genomes to unassembled transcriptomic reads. Details can be found in Table S1. To complement these data, we added new sequence data for another 17 species listed in Table S2. Most of these taxa have well characterized reproductive biology (also listed in Tables S1&2), with some exceptions. For the species we sequenced, we confirmed putative sex chromosomes via coverage difference between male and female sequenced reads where possible. For existing datasets, the least well characterized was the recently described *Coronaproctus castanopsis* Li, Xu & Wu (Monophlebidae). The species description and genome assembly do not report the sex determination system (Zheng et al. 2023; Huang et al. 2024); however, evidence strongly suggests that it is XX/X0. First, a recent molecular phylogeny based on genomic and transcriptomic data confirmed its status as a monophlebid and suggests that *C. castanopsis* shares a more recent common ancestor *Drosicha* spp. (Kuwana) than it does with the hermaphroditic, haplodiploid *Icerya* spp. (Song et al. 2025). *Drosicha* Walker species have been previously reported as XX/X0 (Yokogawa and Yahara 2009) and in synteny analyses we found orthology between the smallest chromosome of *C. castanopsis* and the smallest chromosome of *D. contrahens*, which itself is synthetic to the newly identified X of *Neosteingelia texana*. Additionally, in the haplodiploid hermaphrodites like *Icerya purchasi*, production of pure males is known to be exceedingly rare (e.g., a recent sample of over 300 adults, collected from 27 populations across the world recovered no males: Mongue et al. 2021). The species descriptions for *C*. *castanopsis* include both male and female morphology with no indication that males were rare or difficult to obtain (Li et al. 2023; Zheng et al. 2023), suggesting roughly equal sex ratios expected of chromosomal sex determination.

### Collecting samples from across sex determination systems in scale insects

For newly generated data, we sampled species across sex determination systems of scale insects, with special interests in targeting representatives of families that had no existing sequence data and/or with documented X chromosomes (Gavrilov 2007), but previously karyotyped species were not always readily available for collection. Many of these scale insects have cryptic life cycles and spend much of the year under tree bark or underground (Kosztarab and Watson 1994; Thomson et al. 2021). In several cases, we were able to obtain only a few individuals from a target species and sometimes only females, not enough starting tissue to generate chromosome-level genomes, perform the sex chromosome analyses, or differential expression analyses described below. Still, these samples represent new sequencing resources for an understudied group of insects and help corroborate existing hypotheses of phylogenetic relationships.

We aimed to sequence DNA from a single individual where possible, but for some smaller taxa, we resorted to pooling multiple individuals. We sequenced these samples for inclusion in the larger phylogeny, but given the fragmented nature of their draft genomes, we did not perform the synteny and orthology analyses described below. Species sequenced under this strategy include: *Orthezia annae* Cockerell (Ortheziidae), *Insignorthezia insignis* (Browne) (Ortheziidae), *Neogreenia osmanthus* (Yang & Hu) (Qinococcidae), *Eumargarodes laingi* Jakubski (Margarodidae), *Stomacoccus platani* Ferris (Steingeliidae), *Puto decorosus* McKenzie (Putoidae), *Puto cupressi* (Coleman) (Putoidae), *Nipaecoccus nipae* (Maskell) (Pseudococcidae), *Nipaecoccus floridensis* Beardsley (Pseudococcidae), *Pseudophilippia quaintancii* Cockerell (Coccidae), and *Pityococcus rugulosus* McKenzie (Pityococcidae). Of these, *P. decorosus* was the only species for which we obtained a male. For more information on collected species, please see Table S2.

We were able to find an abundant ensign scale (Ortheziidae): the Spanish moss ensign scale, *Graminorthezia tillandsiae* (Morrison) and a member of Kuwaniidae: the giant pecan scale, *Neosteingelia texana* Morrison. Because much is still unknown about the ecology and life history of these rarely-studied scale insects, we provide brief observational notes on each in the supplement. We also sequenced two hermaphroditic monophlebids, *Crypticerya genistae* (Hempel) and *Crypticerya rileyi* (Cockerell). We followed the same general sequencing strategy for these species, with modifications noted below. Because scale insects are highly sexually dimorphic, including in body size (with females being the larger sex, Vea et al. 2016) and the expected karyotypes are either fully haplodiploid or XX/X0, we based our primary assemblies on females. To identify sex chromosomes, we used a typical coverage-based approach (Mongue et al. 2017, 2022; Mongue and Baird 2024; Baird et al. 2025), and sequenced DNA from pooled males. Slide-mounted voucher specimens associated with each of the aforementioned collections are deposited in the Florida State Collection of Arthropods (FSCA), Gainesville, Florida, USA. We additionally obtained a limited sample of an unidentified *Puto* species: one adult male and five adult females. To make the most of this valuable XX/X0 sample, we chose a distinct sequencing approach, as described below.

### DNA extraction and sequencing approach

For all samples, we used a modified Omniprep (G-Bioscences, St. Louis, MO, USA) chloroform extraction and ethanol precipitation protocol, following previously successful application in scale insects (Mongue et al. 2024; Liesenfelt et al. 2025a). Specifically, we used a dounce homogenizer to grind whole individuals, extended the lysis step overnight, and used 2 µL of 20mg/mL glycogen to facilitate DNA precipitation during a 1hr rest at −20C. In the case of *N. texana* we used a single adult female, but because *G. tillandsiae* is much smaller (length <1mm), we pooled seven females into one extraction. We sent DNA extractions from *N. texana, C. genistae*, and *C. rileyi* to Arizona Genomics Institute (AGI) (Tucson, AZ, USA) for standard PacBio HiFi library prep and sequencing on a Revio (Pacific Biosciences, Menlo Park, CA, USA). We sequenced *G. tillandsiae* with the University of Florida’s Interdisciplinary Center for Biotechnology Research (Gainesville, FL, USA) where we used a low-input library approach and sequenced on a Sequel IIe machine.

To identify the X sex chromosomes in these species, we separately sequenced DNA from males of both species to assess differences between autosomal and sex chromosome coverage. For male extractions, we used the same extraction protocol, but in both species we pooled five individuals per extraction. Once extracted, we sent samples away for sequencing on an Illumina Novaseq at Novogene (Sacramento, CA, USA).

Separately, we snap froze (from live to −80°C) 50mg of female tissue from each species for Hi-C protocols. We sent frozen tissue to AGI for crosslinking and sequencing. For both this and the male sequencing above, we waited until we received initial PacBio data and calculated genome size to ensure we requested adequate coverage (∼50x).

For the *Puto* sp. sample, we extracted DNA from the single male using the protocol above but falling short of the input thresholds for standard HiFi library preparation, we opted to use the PacBio AmpliFi whole-genome amplification kit. We then submitted the female tissue, preserved in ethanol, for Hi-C library preparation with AGI. Thus, although we used a different strategy, ultimately, we still had sequencing from a male to identify the X chromosome by coverage.

### Genome assembly and validation

For all species, we received raw PacBio HiFi reads. We started by passing the reads to jellyfish v2.3.0 (Marcais and Kingsford 2012) to count 21-mers and build a distribution of the frequency of these sequences. Simple calculations on these frequencies gave us both an expectation for genome size and depth of sequencing coverage. Next, we assembled raw reads with hifiasm v0.8 (Cheng et al. 2021). For this step, we included the −l 3 parameter for minimizing duplicate haplotig retention and set the --hom-cov parameter to the peak observed in our k-mer counting step. For these initial assemblies, we calculated basic summary statistics (N50, sequence count, etc.) and ran BUSCO v5.8.3 using the hemiptera_odb10 lineage to assess completeness and duplication (Manni et al. 2021).

For each species, we took the primary assembly through the purge_dups pipeline published only as a git repo (https://github.com/dfguan/purge_dups) to identify cases in which both haplotypes were kept as duplicate sequences in the assembly via a mix of coverage and sequence identity during self-alignment. Note that purging duplicates is an iterative process and can be run in successive rounds. Indeed, because we pooled multiple *G. tillandsiae* as a single sample, we expected retained haplotigs to be a larger issue in its assembly. We assessed BUSCO scores and assembly stats after the first round and ran a second round of duplicate purging on the *G. tillandsiae* assembly based on the high rate of duplication.

After purging, we aligned Hi-C sequencing reads following Arima’s recommended pipeline (detailed on their git repo: https://github.com/ArimaGenomics/mapping_pipeline). In brief, we quality trimmed reads using the repo’s custom scripts, separately mapped each of the two mate pairs to the draft genome using BWA v7.9a (Vasimuddin et al. 2019), and combined alignments with Arima’s scripts. We used the merged alignments as inputs for YaHS v1.1 (Zhou et al. 2023). After scaffolding, we again assessed assembly statistics before manual curation. For this step, we used the YaHS command “juicer pre” to generate inputs for manual review in Juicebox v2.17.00 (Durand et al. 2016). We made minor adjustments based on contacts as needed, output the updated scaffold relationships, and updated the assembly with YaHs “juicer post” command (Zhou et al. 2023). For the final time, we calculated assembly statistics and BUSCO completeness (Manni et al. 2021).

To identify the sex chromosomes, we aligned male short-read sequencing to the respective finalized reference using Bowtie2 v2.5.4 (Langmead and Salzberg 2012) using the “--very-sensitive-local” alignment parameters. We then coordinate-sorted the alignments with Picard tools v3.2.0 (Wysoker et al. 2013) and used SAMtools v1.9’s coverage function (Danecek et al. 2021) to calculate coverage across scaffolds. For the *Puto* sp., we mapped long reads to the reference with minimap2 VERSION (Li 2018) using the “-x map-hifi” parameter to optimize for that data type. Based on the expectation of X0 karyotype for males, the X chromosomal scaffold should have 50% the coverage depth of the diploid autosomes.

### Phylogenomic analysis

We followed the BUSCO_phylogenomics pipeline (https://github.com/jamiemcg/BUSCO_phylogenomics). For each genome or transcriptome included in the analysis, we assessed the BUSCO completeness as above (again using the hemiptera_odb10 dataset). Then, using the scripts provided by the git repo, we parsed single-copy BUSCO amino acid sequences, with a threshold of presence in 75% of samples for inclusion. In the analysis presented here, that totaled to 1,477 single-copy protein sequences. The pipeline produced trimmed protein alignments and individual gene trees, which we analyzed in two ways. First, we concatenated the individual gene alignments with custom bash scripts and used IQTree (Minh et al. 2020) v2.2.2.7 to compute the maximum likelihood tree using the “Q.insect+FO+G8” model of substitution, with confidence assessed with 1000 ultrafast bootstraps and Shimodaira-Hasegawa-like approximate likelihood ratio test (SH-aLRT) (Guindon et al. 2010).

Second, to offer an additional phylogenetic hypothesis that accounts for incomplete lineage sorting, we used ASTRAL (Zhang et al. 2018) to generate a consensus species tree from the individual gene trees with confidence assessed as posterior quartet probabilities (Sayyari and Mirarab 2016). As both trees showed agreement, but the species tree approach offers less informative branch lengths, we present only the maximum likelihood approach in the main text and the ASTRAL approach in the supplement. In both cases, we manually rooted trees with *Acyrthosiphon pisum* (Harris) and *Adelges tsugae* (Annand) as the non-scale outgroups, visualized the output with FigTree v1.4.4 (Rambaut 2010), and manually colored branches by sex determination type.

### Annotation and macrosynteny analysis

With finalized assemblies, we next generated gene annotations for downstream analyses. First, we used RepeatModeler v2.0 (Flynn et al. 2020) to *de novo* identify repetitive sequences, including the --LTR parameter to search for long terminal repeats. We passed the identified repeats to RepeatMasker v4.0.9 (Smit et al. 2019) to generate a softmasked assembly ready for annotation. We used a standard approach across species: BRAKER v3.0.8 (Hoff et al. 2019) with protein evidence from collected BUSCO protein sequences across scale insects. To minimize biases in downstream analyses arising from different annotation pipelines, we re-annotated published assemblies with the BRAKER protein pipeline described above Then, we used MCScanX v1.0.0 to infer synteny (Wang et al. 2024). This method uses BLAST similarity to identify putative 1-to-1 orthologs between pairs of species and then takes advantage of the scaffold coordinate information in gene annotations to identify regions of the genome in one species that correspond to the orthologous region of the other. We based pairwise comparisons on the phylogenetic relationships inferred above.

### Conservation of X-linked genes

Once we established the identity of the X chromosomes, we examined in more detail the fate of X-linked genes across the losses of the X in hermaphrodite and the paternal genome elimination clades. In particular, we examined two patterns. First we subsetted our gene annotations for *G. tillandsia*, *N. texana*, *C. castanopsis*, and *Puto* sp. to only compare genes on the X chromosomal scaffolds of these species to each other and to the full genomes of either (1) *I. purchasi*, *C. genistae,* and *C. rileyi* or (2) the chromosome level assemblies of the PGE species to look for conserved, X-linked genes lost in the transition to other sex determination systems. Second, we compared the full genomes of the X sex determination species to the full genomes of groups (1) or (2) above to look for any genes conserved in the hermaphrodites or paternal genome elimination species but missing from the X species. We used OrthoFinder v2.5.2 (Emms and Kelly 2019) to identify orthogroups. We then searched NBCI’s RefSeq protein database to infer functional information via sequence homology.

### Expression data generation and analysis

We sought to further characterize the state of the X chromosome both before and after its loss by examining patterns of gene expression between males and females. In practice, we were only able to obtain sufficient samples of both sexes for two species: *N. texana* and *T. liriodendri*. For each of these species, we generated Illumina paired end RNAseq from four biological replicates of individual adult females and four replicates of pooled males. For *N. texana*, we used pooled replicates of 2–4 males per extraction and for *T. liriodendri* we used replicates of 5 males per extraction.

For RNA extractions, we utilized a RiboPure™ RNA Purification Kit (Invitrogen™) with slight modification to the protocol to account for low tissue mass of our samples. We thawed previously frozen tissue directly into 500 µL of the kit provided TRI reagent to account for tissue mass < 50 mg. With a lower starting volume, the aqueous phase measured ∼ 250 µL after phase separation. To keep the 1:2 volume ratios consistent to those in the protocol, we added 125 µL of 100% ethanol to this aqueous phase for RNA purification. Extractions were then eluted into 50 µL of buffer instead of the recommended 100 µL. Our extractions were finally sent to Genewiz (South Plainfield, NJ, USA) to generate 30 million paired end Illumina reads per sample.

Added to these newly sequenced samples, we downloaded expression data for one adult male and two early adult female *Ericerus pela* (Chavannes) from the NCBI Sequence Read Archive (SRA) (accessions:SRR1027692 and SRR9617905 + SRR9617914, respectively). For each SRA accession, we quantified per-sample expression against the BRAKER annotation generated above using Kallisto v0.51.1 (Bray et al. 2016). We analyzed expression as TPM (transcripts per million) using R scripts to calculate SPM, the specificity metric, (Kryuchkova-Mostacci and Robinson-Rechavi 2017), as the proportion of each gene’s expression in one sex compared to the other. We chose this approach as a calculated tradeoff to make the most of available data, as the single male sample from *E. pela* would not allow for more traditional differential expression analyses, at the cost of losing statistical evaluation of expression difference. We plotted sex specificity as the amount of expression in females, a value ranging from zero (male-limited expression) to one (female-limited expression). Additionally, following previous evolutionary genetic analyses (Mongue et al. 2022), we binned genes into categories of male-biased (SPM < 0.3), unbiased (0.3 < SPM < 0.7), or female-biased (SPM > 0.7). With these categories, we used X^2^ tests of independence to assess whether the sex-biased composition of genes different between X or former-X and autosomes in *N. texana*, *E. pela*, and *T. liriodendri*.

Finally, we explored dosage compensation in the one X chromosome species for which we have RNA data, *N. texana*. In particular, we took the subset of genes expressed only in males or only in females and calculated the mean of each gene’s TPM expression across four biological replicates for that sex and compared it across autosomal and X-linked genes using a Wilcoxon test. Because we found a majority of genes with sex-limited expression in our SPM analyses, we did not directly compare expression between the sexes.

## Results

### Assemblies, X assignments, and annotations

We assembled 17 new scale insect genomes to various levels from basic Illumina-only unordered contigs to Hi-C scaffolded chromosome level assemblies. For brevity and to keep the focus on the evolutionary genomic findings, we describe the full assembly statistics and report accessions in the supplement. We used assemblies of all levels for the phylogeny to infer turnover of sex determination, but proceeded to further analysis of synteny, orthology, and gene expression for only the subset of chromosome-level assemblies.

### An updated phylogeny of scale insects

We reconstructed the phylogeny of scale insects using a mix of publicly available and newly generated datasets. We used two different methodologies, a maximum likelihood inference of sequence evolution using concatenated protein sequences and a consensus species tree inferred from individual gene trees. We present the maximum likelihood approach in the main text (**Figure 1**) and the consensus species tree in the supplement (**Figure S1**). Both approaches yielded the same topology and had high support metrics for relationships;therefore, the following results summary applies to both analyses.

**Figure 1.**
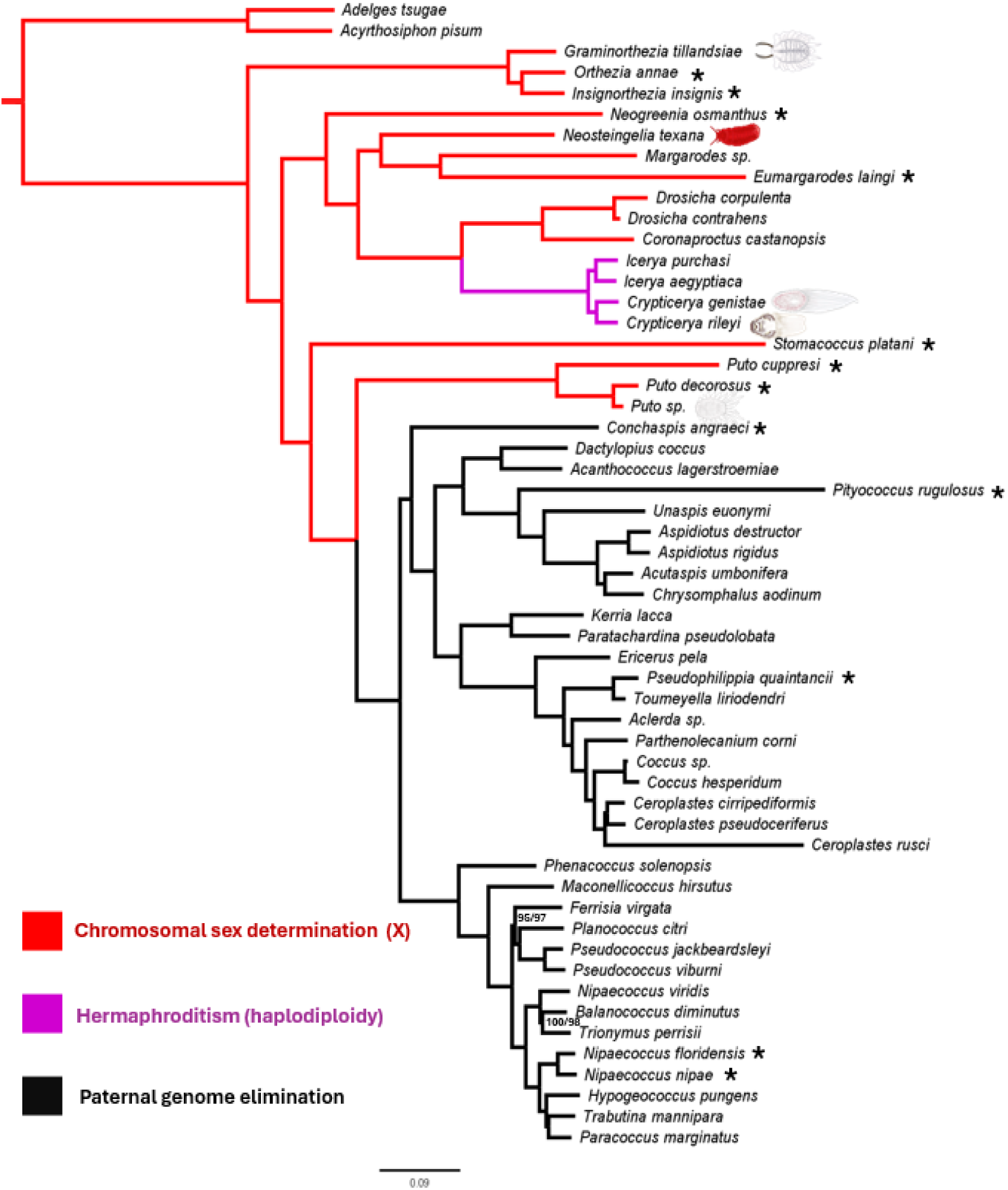
Maximum likelihood phylogeny of scale insects annotated with sex determination system. Relationships inferred from concatenated BUSCO amino acid sequences from genomes and transcriptomes. Support values follow the format Shimodaira-Hasegawa-like approximate likelihood ratio test (SH-aLRT) / Ultrafast Bootstrap values; all nodes have 100/100 support unless otherwise noted. Sex determination systems are color-coded as follows: red - X chromosomal sex determination, purple - haplodiploid hermaphroditism, black - paternal genome elimination. Species with an * next to their label have newly sequenced draft genomes, those with illustrations have newly sequenced chromosome-level assemblies for this study. Branch lengths are proportional to the expected number of amino acid changes per site as indicated by the scale bar.

We rooted the tree with known outgroups, the pea aphid, *Acyrthosiphon pisum* and hemlock woolly adelgid, *Adelges tsugae* (Johnson et al. 2018). We recovered most relationships in agreement with other published phylogenies (Gullan and Cook 2007a; Cook et al. 2002; Hodgson and Hardy 2013; Vea and Grimaldi 2016), so we focus on the main differences and additional insight gained from newly sequenced taxa here. First, we recovered Ortheziidae as a monophyletic group and the earliest diverging family of scale insects. In fact, based on current sampling, all other family level groupings are valid with the exception of Coccidae. Our sole representative, a non-specified member of the genus *Aclerda*, is placed confidently in the middle of Coccidae. Aclerdids could represent a lineage of grass-specialist soft scales (though not all aclerdids are found on grasses, see García Morales et al. 2016 for a review) and its validity as a family may need to be reevaluated. Putoidae is recovered as a valid family falling out between what has historically been considered ‘Archaeococcoidea’ and ‘Neococcoidea’.

However, we do not find evidence that these two superfamily groupings are monophyletic. We lack resolution to validate most genus-level relationships, but members of *Ceroplastes*, *Coccus*, *Aspidiotus*, *Pseudococcus*, *Icerya*, *Crypticerya*, and *Puto* all share a more recent common ancestor with a congener than any other taxon. The one exception is the genus *Nipaecoccus*; *Nipaecoccus floridensis* and *N. nipae* group together as expected, but *N. viridis* is confidently placed closer to *Balanococcus diminutus* than to other *Nipaecoccus*, suggesting the generic placement of *N*. *viridis* may need to be revisited.

Finally, most importantly for the purposes of this study, the relationships described above, combined with existing knowledge of family level reproductive systems (reviewed in Gavrilov 2007), suggests a straightforward scenario for sex determination turnover. Parsimoniously, X chromosomal sex determination was lost twice, once in the lineage of simultaneous hermaphrodites within Monophlebidae (here represented by *Icerya* and *Crypticerya*), and a second time independently at the base of the group of paternal genome elimination families including Pseudococcidae, Coccidae, Diaspididae, and others. We also explored the evolution of different types of paternal genome elimination, but lacking information for most sequenced Coccidae species, we cannot infer the timing or number of transitions within paternal genome elimination. The only straightforward inference, based on the state of early-diverging taxa, is that lecanoid paternal genome elimination appears to have evolved first, with Comstockiella and diaspidid types arising secondarily (Figure S2).

### Macrosynteny: two divergent paths to losing X sex determination

We mapped 1-to-1 orthologs between species across the subset for which we have chromosome-level assemblies. First, we consider the Monophlebidae (Figure 2). Starting with the outgroups, despite the X having split into two chromosomes, an X_1_X_2_ system, in *Adelges tsugae*, both of these sex chromosomes are syntenic to the single X chromosome of *Acyrthosiphon pisum*, thus the X is conserved in the outgroups. However, we found overall very little synteny between our earliest diverging scale insect, *G. tillandsiae*, and other Sternorrhyncha. Interestingly, what little detectable orthology exists between the *G. tillandsiae* and *Acyrthosiphon pisum* is autosome-to-autosome, suggesting that the X chromosome has highly diverged between the scale insects and other Sternorrhyncha, or potentially even turned over.

**Figure 2.**
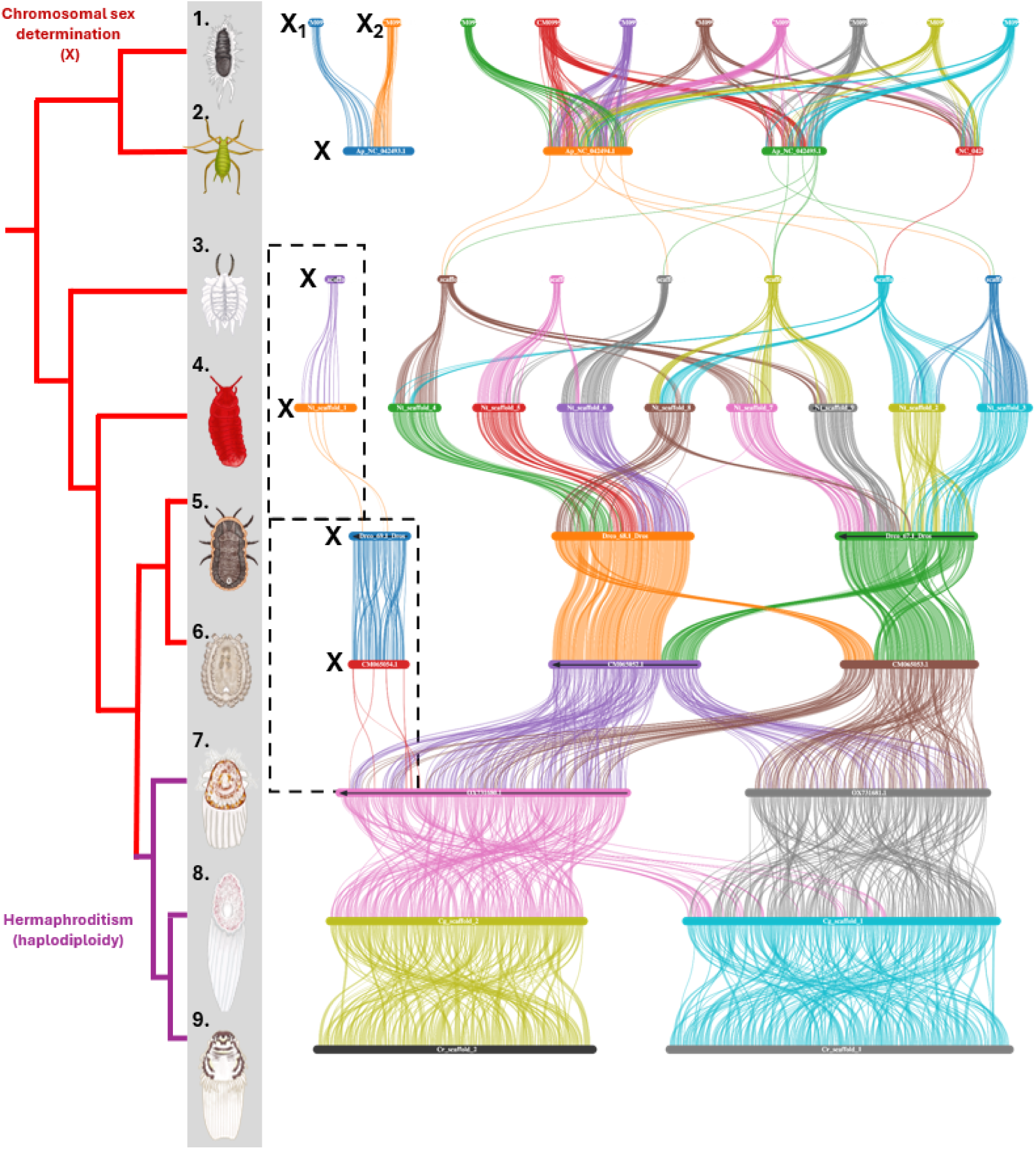
Macrosynteny across the transition from X to simultaneous hermaphroditism. Individual colored horizontal bars represent chromosomal scaffolds. Lines connecting them represent conserved orthologs between a pair of species. For each species, where applicable, the X chromosome(s) are sorted to the left-most side of the plot and labeled. The dashed box highlights the fate of X-linked orthologs across the turnover in sex determination systems.

Phylogenetic relationships follow those recovered in Figure 1 with the same coloring for branches. Species depicted: 1. *Adelges tsugae* 2. *Acyrthosiphon pisum* 3. *Graminorthezia tillandsiae* 4. *Neosteingelia texana* 5. *Drosicha contrahens* 6. *Coronaproctus castanopsis* 7. *Icerya purchasi* 8. *Crypticerya genistae* 9. *Crypticerya rileyi*. Illustrations courtesy TL.

Next, comparing *G. tillandsiae* to *N. texana*, we found clearer evidence for a conservation of the X chromosome. Both species’ X chromosomes were confirmed by coverage difference between males and females within species (see Supplement), and the two chromosomes only share orthologous hits with each other. There are some rearrangements to the karyotype, as expected when comparing n = 9 vs. n = 7 chromosomes (based on Hi-C contacts) but overall much more conservation of orthologs.

Regarding, *N. texana* and *D. contrahens*, we again found good evidence for conservation of the X chromosomes, but note that the assembly of *D. contrahens* contains no sex linkage information. This genus is known to be XX/X0 based on karyotype, thus we inferred the identity of the X via synteny to the *N. texana* X. In contrast to the conserved X, we found substantial reorganization of the autosomes. *Drosicha contrahens* has a mere n = 2 autosomes, which appear to have resulted from simple fusions given that these two autosomes map cleanly to 3 and 4 of the *N. texana* autosomes, with only 1 inferred to have orthology to multiple *D. contrahens* autosomes. Intriguingly, this suggests that the extreme karyotype reduction found in the largely asexual *Icerya* and *Crypticerya* (n = 2) began *before* the transition to asexuality, not as a consequence of it.

Considering *D. contrahens* and *C. castanopsis*, we recovered additional evidence that the latter is an XX/X0 sex determination system. Namely, the smallest of the three chromosomes (CM065054.1) shows fewer orthologs than the other chromosomes and only has hits to the X of *D. contrahens*. There appears to have been autosomal rearrangement between lineages here, as one end of each autosome maps to the opposite autosome in the other species. Given that we recover the same pattern between *C. castanopsis* and *I. purchasi* below though, it is more parsimonious to conclude that this rearrangement happened once in the lineage leading to *Coronaproctus* rather than shuffling and and reverting between multiple lineages.

Finally, we recovered much more shuffling in the comparison that spans the loss of X sex determination between *C. castanopsis* and *I. purchasi*. The little remaining orthology to the X maps to the larger of the two autosomes, which seems indicative of an X-autosome fusion, but this only captures a tiny fraction of the rearrangements. The two autosomes of *C. castanopsis* do not cleanly correspond to the two autosomes of *I. purchasi*. The two large inter-chromosomal rearrangements may reflect a derived karyotype in *Coronaproctus*, as mentioned above, but we also recovered myriad intra-chromosomal rearrangements within the autosomes between these species. Continuing further into the *Crypticerya* species, we found additional rearrangements both within and between chromosomes with plenty of conserved genes but very short blocks of synteny.

For the second loss of X sex determination, we explored the transition to paternal genome elimination (Figure 3). First, we confirmed that the X of the closest sequenced outgroup to paternal genome elimination, *Puto* sp., is syntenic to the scale insect X via comparison to *N. texana.* Next, we traced the orthologous chromosome into the paternal genome elimination clade and found, surprisingly, that it is essentially intact, albeit no longer sex-determining, in the crapemyrtle bark scale, *Acanthococcus lagerstroemiae* (Kuwana), where it is orthologous to chromosome 7 in the most complete assembly (scaffold CM063112.1). Indeed, this chromosome is well conserved in *Kerria lacca* (Kerr) and *Ericerus pela* and only begins to break apart as part of a broad, substantial increase in chromosome number in *Toumeyella liriodendri* (Gmelin) and *Ceroplastes pseudoceriferus* Green (n = 17 and 18, respectively). These karyotype changes appear to be simple fissions as most *E. pela* chromosomes correspond to two *T. liriodendri* chromosomes with an additional translocation from an autosome. In *Ce. pseudoceriferus*, the former X has split into three smaller chromosomes. Oddly, the closest sequenced relative of *Ce. pseudoceriferus*, *Coccus hesperidum* (Linnaeus), has a reduced karyotype (n = 7) that appears to have resulted from secondary fusion of the fragmented chromosomes as the fewer chromosomes here incorporate similar, but sometimes distinct, syntenic blocks compared to the *E. pela* genome. In this species the remnant X has fused with a syntenic block that was previously autosomal in all other sequenced species. *Coccus hesperidum* also marks the last point at which the former X is easily distinguishable in the paternal genome elimination species.

**Figure 3.**
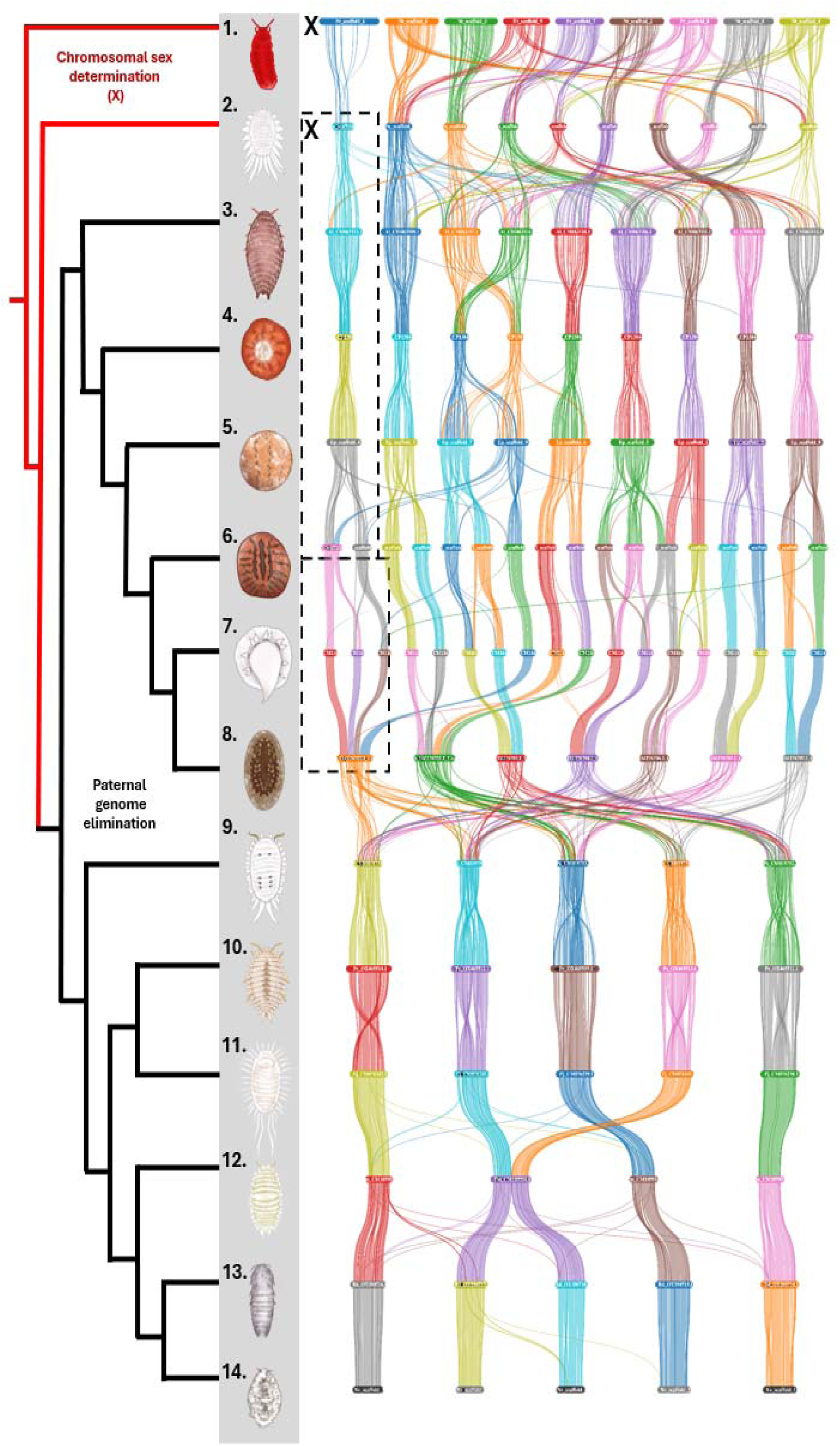
Macrosynteny across the transition from X to paternal genome elimination. Individual horizontal bars represent chromosomal scaffolds. Lines connecting them represent conserved orthologs between a pair of species. For each species, where applicable, the X chromosome is sorted to the left-most side of the plot and labeled. The dashed box highlights the fate of X-linked orthologs across the turnover in sex determination systems. Phylogenetic relationships follow those recovered in Figure 1 with the same coloring of branches. Species depicted: 1. *Neosteingelia texana* 2. *Puto* sp. 3. *Acanthococcus lagerstroemiae* 4. *Kerria lacca* 5. *Ericerus pela* 6. *Toumeyella liriodendri* 7. *Ceroplastes pseudoceriferus* 8. *Coccus hesperidum* 9. *Phenacoccus solenopsis* 10. *Planococcus citri* 11. *Pseudococcus jackbeardsleyi* 12. *Paracoccus marginatus* 13. *Balanococcus diminutus* 14. *Nipaecoccus viridis*.

There have been major rearrangements across all chromosomes leading to the true mealybugs (Pseudococcidae) and formerly X-linked orthologs are scattered across all five chromosomes of *Phenacoccus solenopsis* Tinsley. After this rearrangement, however, genome structure is remarkably well conserved. Five of the six chromosome-level assemblies show the n = 5 karyotype, with only individual-level gene-trafficking events rearranging synteny for the most part, barring two exceptions. First, *Paracoccus marginatus* has only n = 4 chromosomes, though this reduction is the result of a single, simple fusion of the second and fourth linkage groups in other mealybugs. Second, even microsynteny (*i.e.*, the individual ordering of orthologous genes within chromosomes) is well preserved across most species, but in comparisons between *Ph. solenopsis* and *Planococcus citri* (Risso), as well as *Pl. citri* and *Pseudococcus jackbeardsleyi* Gimpel & Miller, we recovered large inversions on four of the five chromosomes.

### Conservation of genes across sex determination turnover

We searched for orthologs with two distinct conservation patterns: first, those conserved on the X chromosomes but absent from the non X-chromosomal scales and, second, those absent from anywhere in the genome of X sex determination species but conserved across all hermaphrodites or paternal genome elimination species.

Starting with the former, we searched across the X chromosomes of *G. tillandsiae* (n = 3,534 genes), *N. texana* (2,497 genes), *C. castanopsis* (1,967 genes), and *Puto* sp. (1,633) compared to the entire genomes of *I. purchasi* (18,371 genes), *C. genistae* (20,330), and *C. rileyi* (19,871 genes). We recovered 410 orthogroups conserved across all the X chromosomes. Of these conserved X-linked orthologs, only 8 were absent from all hermaphroditic species (**Table S5**).

Similarly, but even more extreme for the paternal genome elimination clade, we identified only two conserved X-linked orthologs that were missing from all sampled paternal genome elimination species (**Table S6**).

We also searched for the inverse pattern, novel gene gain in the hermaphrodites or paternal genome elimination species. We found 664 orthogroups conserved in all three hermaphrodite species that were missing from all of our annotated X sex determination species. A majority of these (449, 68%) had no significant BLAST hits and a further 54 matched to “hypothetical” or “uncharacterized” proteins, leaving 161 with functional hits. There are too many to parse here, and none appear as immediately crucial to sex determination, but all of these orthogroups, as well as their top BLAST hits are listed in **Tables S5–6**.

Turning to paternal genome elimination, we found 49 orthogroups that were gained and conserved across all sampled paternal genome elimination species. Of these, 18 (37%) returned “hypothetical protein” hits or no significant matches. Given the unique nature of paternal genome elimination, its molecular functions may lie in these poorly characterized proteins, but we can do no more with this information at present. We identified one better characterized orthogroup as a DDB1- and CUL4-associated factor 8, which has been implicated in cancer in humans via changes in methylation regulation (Yang et al. 2015). Less directly related to reproduction, but interesting nonetheless, we also identified an orthogroup related to fatty acyl-CoA reductase, which is conserved as a single copy ortholog in *A. lagerstroemiae* and *K. lacca*, but in multiple copies, from 2 to 16, in other paternal genome elimination species. This apparent gene family expansion may be related to the diversity of wax production across the group (Foldi 1991; Ding et al. 2022). These genes and BLAST hits are cataloged in **Table S7.**

### Patterns of gene expression and sex-biased genes before and after a loss of X

Finally, we generated RNA sequencing from whole male and female samples of *N. texana* and *T. liriodendri*, downloaded public data for *E. pela*, and analyzed patterns of sex-specific gene expression. First, we explored gene expression levels, as measured by TPM (transcripts per million), between the X and autosomes of *N. texana* with males and females separately (Figure 4**, left**). As expected, expression levels did not differ between the autosomes and diploid X of females (W = 61230, p-value = 0.880). The haploid X in males also expressed genes at a comparable level to the autosomes; in fact, expression trended higher for X-linked genes here (W = 29963, p-value = 0.060). As such, it appears dosage of the haploid X is upregulated in females. Owing to extremely dimorphic gene expression and few genes expressed roughly equally between the sexes, we did not directly test expression levels between sexes.

**Figure 4.**
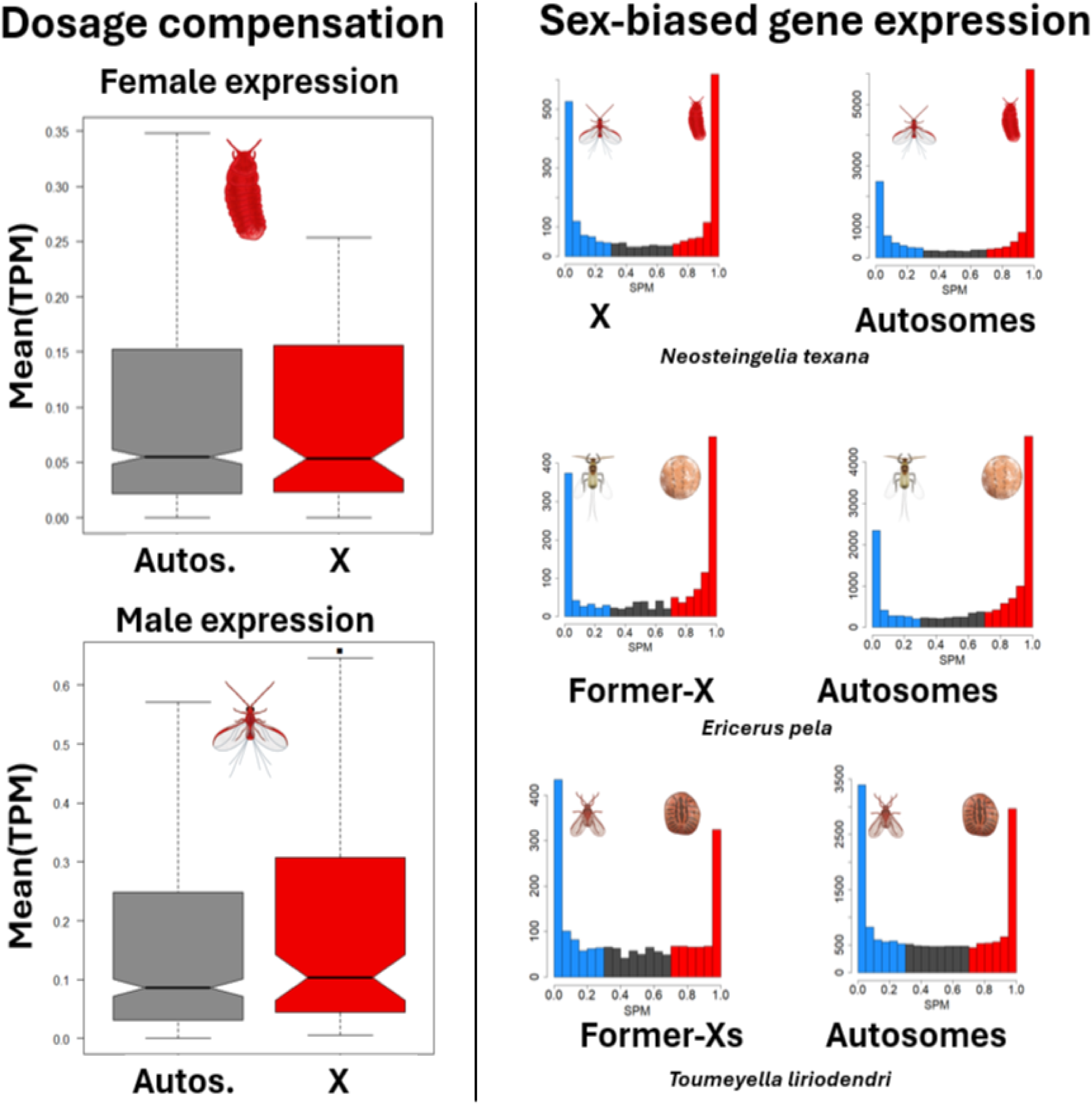
Gene expression patterns within and between sexes. Left: Dosage compensation in *Neosteingelia texana*. Genes on the X and autosomes show similar expression levels (**top**) in females as well as (**bottom**) in males, as expected if the haploid X is compensated to express genes at a higher level in males.The dot represents a trend for higher expression on the male X compared to autosomes. **Right: Patterns of sex-biased gene expression across species.** In each row, individual plots show the SPM (specificity metric) of expression in male vs. female tissue. In each case, lower values represent male-biased expression, with SPM = 0 being male-limited expression; SPM = 1 female-limited expression.The **left** histogram shows the distribution of expression patterns for the X or its orthologous chromosome(s) after the transition to paternal genome elimination. The **right** histogram shows the same for (all other) autosomes.

Second, we explored the distribution of genes with sex-biased expression. Starting with *N. texana*, surprisingly, we found that while the X still has a plurality of female-biased genes, it holds proportionally fewer than the autosomes, as well as proportionally more male-biased genes (X^2^_2_ = 93.15, p < 0.0001, Figure 4**, right, top; Table 1**). Even after the loss of the sex determining function in *E. pela*, the former-X chromosome still has a different composition than the autosomes (X^2^_2_ = 100.13, p < 0.0001, Figure 4**, right, middle; Table 1**), with fewer female-biased and more male-biased and unexpressed genes. In *T. liriodendri*, which has had both fissions and fusions of the former-X, there is no difference in the proportions of sex-biased genes between the former-Xs and autosomes (X^2^_2_ = 1.91, p = 0.591, Figure 4**, right, bottom; Table 1**).

**Table 1.** Distribution of sex-biased genes across the genomes of X and paternal genome elimination species. Numbers represent raw counts of genes by category, parentheses give the percentage of the total genes in that group. Bolding highlights significant differences between linkage categories for *N. texana* and *E. pela*.

| Species | Linkage | Male-biased | Unbiased | Female-biased | No expression |
| --- | --- | --- | --- | --- | --- |
| *Neosteingelia texana* | **X** | **864 (37%)** | **301 (13%)** | **971 (42%)** | **194 (8%)** |
|  | Autosomes | 4,651 (29%) | 1,849 (11%) | 8,544 (52%) | 1,323 (8%) |
| *Ericerus pela* | **Former-X** | **513 (24%)** | **226 (10%)** | **808 (38%)** | **3401 (28%)** |
|  | Autosomes | 3,637 (21%) | 2,151 (13%) | 7,847 (46%) | 599 (20%) |
| *Toumeyella liriodendri* | Former-Xs | 772 (34%) | 449 (20%) | 691 (30%) | 365 (16%) |
|  | Autosomes | 6,282 (33%) | 3,878 (20%) | 5,882 (31%) | 2,900 (16%) |

## Discussion

We studied the turnover of reproductive systems across scale insects. To do so, we generated new chromosome-level references for the Spanish moss ensign scale (*Graminorthezia tillandsiae*), the giant pecan scale (*Neosteingelia texana*), a giant mealybug (*Puto* sp.), as well as two giant scales (*Crypticerya genistae* and *C. rileyi*) in service of understanding the changes to genome architecture associated with change in sex determination. However, before considering sex determination further, these new genomic resources offer a chance to update and confirm phylogenetic relationships across a taxonomic group that had, until recently, been represented by a dearth of genomic data.

### Insights from a new phylogeny of scale insects

Overall, our results agree with previous findings, with a few notable exceptions and additions. First, the largest mainly morphological phylogeny includes an early split in scale insects between Neococcoidea (containing true mealybugs and allies) and a clade including Monophlebidae, Kuwaniiade, and Ortheziidae (Vea and Grimaldi 2016). A more recent genomic and transcriptomic study included only monophlebid members of the latter group, and thus could not evaluate this relationship (Song et al. 2025). With our sampling, we confidently place Ortheziidae as the earliest diverging family of sequenced scale insects and place Kuwaniidae sister to Monophlebidae among represented families. Our tree topology recovers the group of Ortheziidae, Monophlebidae, and Kuwaniidae (sometimes grouped as part of Archaeococcoidea (e.g., Foldi 2009; Gavrilov-Zimin 2018) as a paraphyletic group.

Putoidae is recovered as monophyletic falling out between what has been considered the ‘Archaeococcoidea’ and ‘Neococcoidea’ groups. Moreover, Putoidae is confirmed to have an X chromosomal sex determination, further supporting morphological and molecular evidence that they should not be placed within PGE Pseudococcide (Downie and Gullan 2004; Hardy et al. 2008; Vea and Grimaldi 2016; Williams et al. 2011; Choi and Lee 2022; Powell and Miller 2025) despite contested historical placement (McKenzie 1967; Gavrilov-Zimin and Danzig 2012; Danzig and Gavrilov-Zimin 2014). We also place the recently described *Neogreenia osmanthus* (Qinococcidae, Zheng and Wu 2023) as the sister taxon to Kuwaniidae, Margarodidae, and Monophlebidae. Despite superficial similarities to and previous placement within Kuwaniidae (Wu and Nan 2012), we recover the ground pearls, Margarodidae, as more closely related to Kuwaniidae than Qinococcidae is. We also sequenced and placed *Pityococcus rugulosus*, the only sequenced member of Pityococcidae, which has been the subject of some disagreement previously. One study placed them as sister to Margarodidae (Gullan and Cook 2007), another as sister to Steingeliidae (in our tree, *Stomacoccus*, Hodgson and Hardy 2013), and yet another placed them close to Diaspididae (Vea and Grimaldi 2016), the armored scales. Our analyses agree with the latter placement. While there is no documentation of Pityococcidae’s genetic system, this placement suggests it employs paternal genome elimination. Cytogenetics could easily confirm or refute this prediction, but our samples were not suitable for these analyses.

Clearly, there is more work to be done on scale insects, especially the early-diverging and species-poor families; within the scope of this study, however, our phylogenetic tree resolves the history of sex determination turnover. In our new phylogeny, the most parsimonious scenario for sex determination turnover is quite straightforward.

Studies have shown that the X sex chromosome has been well-conserved across the order Hemiptera (Pal and Vicoso 2015). Even in the suborder Sternorrhyncha, the group containing scale insects, the X chromosome is conserved across Aphididae, Adelgidae, and Phylloxeridae (Li et al. 2023). Particularly in Aphididae, the X chromosome has been preserved despite extensive autosomal rearrangements (Huang et al. 2025). In contrast, we found very little orthology between the scale insect X and that of non-scale insect Sternorrhyncha. In fact, we failed to recover any blocks of synteny long enough to be visible between *G. tillandsiae* and *Acyrthosiphon pisum*; the only detectable synteny from the X of *G. tillandsiae* mapped to autosomes of the aphid. This lack of conservation raises the possibility that the X chromosome has turned over between lineages. Certainly, there has been enough time, as Vea and Grimaldi’s (2016) phylogeny used fossil calibration to date nodes and placed the split between scale insects and other Sternorrhyncha in the Permian.

Either way, within the scale insects, the X chromosome has been largely conserved between X sex determination species, with the exception of *G. tillandsiae*. Our data suggest there has been an X-autosome fusion in the lineage leading to this species, as it is syntenic to both the X of *N. texana* and one of the autosomes. Consistent with this scenario, we note that Brown et al. observed another ortheziid, *Praelongorthezia praelonga*, had the same diploid chromosome count between males and females (1958), which could occur in an X system with a neo-Y remnant from the X-autosome fusion.

More broadly, we recover two losses of X chromosomal sex determination as the simplest scenario: once in the family Monophlebidae and again in the group leading to the paternal genome elimination clade. In fact, this scenario also holds as most parsimonious in the other recent phylogenies (Vea and Grimaldi 2016; Song et al. 2025), even with the slight differences in tree topology and taxon sampling, so this finding is quite robust. What is novel is the ability to track genetic and genomic changes across these transitions, which stand in contrast to better studied turnover events, in which one sex chromosome is supplanted for another (Vicoso and Bachtrog 2015; Jeffries et al. 2018). In both scale insect turnovers, the X sex chromosome has been replaced by genome-wide haplodiploidy, but in very different forms, as we explore below.

### Leaving X for hermaphroditism

Starting with the Monophlebidae, some members of this group still employ X chromosomes, such as *Drosicha* spp. (Yokogawa and Yahara 2009) and, most likely, *C. castanopsis*. The others in the tribe Iceryini, represented by the genera *Icerya* and *Crypticerya* in our analyses, are fully diploid simultaneous hermaphrodites (Hughes-Schrader 1925; HughesLJSchrader 1963; Kokilamani et al. 2014). These individuals are phenotypically female in all ways except for the presence of ancillary sperm-producing tissue in their gonads (Johnston 1912). It has been proposed that the sperm producing cells are actually a distinct, self-perpetuating cell lineage (Royer 1975); if true, it is perhaps more correct to think of hermaphrodites as individuals with a diploid somatic and ovarian female genotype with a distinct, selfish sperm cell line. Adding to the complexity, in the best studied species, *Icerya purchasi*, rare pure males have also been observed (Royer and Delavaul 1974; Royer 1975; Kim et al. 2011). There is evidence that these males, when present, still mate with hermaphrodites and genetically contribute to offspring (Mongue et al. 2021), making the species an androdioecious reproductive system (Pannell 2002). At present it is unclear how common androdioecy is among the hermaphroditic Monophlebidae (Mongue et al. 2021), but, uniqueness of this system notwithstanding, the most directly pertinent feature is that males are fully haploid (Hughes-Schrader 1925; Kokilamani et al. 2014). So, at the genomic level, sex determination (hermaphrodite or male) presents as genome-wide haplodiploidy.

This transition to this lifestyle is associated with a reduction in karyotype, as all hermaphroditic species studied so far have a mere n = 2 autosomes (Hughes-Schrader 1925; HughesLJSchrader 1963; Hughes-Schrader and Monahan 1966). Based on our analyses, this reduction karyotype actually precedes the change in sex determination, as the more distantly related *N. texana* has n = X + 8 autosomes and the closer relatives to hermaphrodites, *C. castanopsis* and *D. contrahens*, have n = X + 2 autosomes. Despite the large difference in chromosome count, the pattern of rearrangement appears to be relatively straightforward. The X is conserved between the *N. texana* and *D. contrahens*, while seven of the eight autosomes of *N. texana* essentially map entirely to one of the two autosomes of *D. contrahens*, suggesting whole-chromosome fusion events explain most of the difference. Between *D. contrahens* and *C. castanopsis* the X is still conserved but there has been some additional autosomal rearrangement, namely a swap between the ends of the two autosomes. The rearrangements that span the loss of X sex determination between *C. castanopsis* and *I. purchasi* are not nearly as clean, however.

The X chromosome shows orthologous hits only to a portion of the larger of the two *I. purchasi* autosomes but these appear scattered between autosomal orthologs on that chromosome. Genes from two autosomes of *C. castanopsis* have been rearranged both within and between the autosomes of *I. purchasi*. Other asexual lineages are known to exhibit elevated rates of genome rearrangements, often enabled by mobilization of transposable elements (Ferreira de Carvalho et al. 2016), and such a scenario may have occurred here as well. Indeed between the *I. purchasi* and the two *Crypticerya* species, rearrangements both within and between chromosomes have continued at an elevated rate and these genomes are large (>1Gb) and have the high proportions of repetitive sequence masked in our annotation analyses. That said, the sexually-producing *N. texana* had the highest proportion of repetitive sequence in our analysis and a ∼950Mb genome, so large, repeat-rich genomes also predate the transition to hermaphrodites.

To get a more granular sense of changes associated with sex determination turnover, we also performed orthology analyses, searching for conserved X-linked orthologs that were lost from the genomes of the hermaphrodites, as well as genes conserved in the hermaphrodites but missing from the genomes of the X sex determination species. Starting with the former, we found only eight X-linked orthologs missing from all three hermaphrodite species, including: a serine protease stubble-like, a digestive cysteine proteinase-like, a tumor protein p63 regulated gene-like ortholog, and an ortholog of CCN family member 1. Stubble proteins are hypothesized to be secreted signal proteins in another true bug (Bao et al. 2014); cysteine proteinases canonically play roles in metabolism (Goulet et al. 2008), and p63 related genes have a suite of functions from histone interaction to senescence (Sadu Murari et al. 2025). Proteins in the CCN family have functions in neuronal development and gastrulation in vertebrate models (Katsube et al. 2009), but are less well characterized in invertebrates (Hu et al. 2019). In summary, none of these is an obvious candidate for sex determination. It could be that they perform non-canonical functions in scale insects, but it is also possible that sex determination turnover has occurred in a similar way to that proposed for like-for-like turnovers, through addition rather than subtraction of genetic elements (van Doorn and Kirkpatrick 2007). Indeed, we found over 600 orthogroups conserved across the hermaphrodites but missing from the X sex determination species. A majority of these had no database hits, as expected of true novelty. This number could be inflated relative to the paternal genome comparison below for the simple reason that we searched for conservation across fewer hermaphrodite species than paternal genome elimination species, but it is also consistent with the overall more dramatic change during sex determination turnover here. Further sequencing of Monophlebidae can hopefully narrow down a more tractable set of candidate genes for more extensive study.

### Leaving X for paternal genome elimination

The second loss of X chromosomal sex determination gave rise to the paternal genome elimination clade of scale insects. Based on our recovered phylogenetic relationships, the different forms of paternal genome elimination found across scale insects (reviewed in Gavrilov 2007) represent elaborations from a single origin rather than independent evolutions. Based on data from early-diverging paternal genome elimination species, we infer that the ancestral type of paternal genome elimination is lecanoid, in which males keep but do not express paternal chromosomes in the soma (de la Filia et al. 2021) and only truly eliminate them during spermatogenesis (Hughes-Schrader 1948; Nur 1980). This sequence of paternal genome elimination evolution makes intuitive sense. All types of scale insect paternal genome elimination are united by haploid male expression and elimination of paternal chromosomes from sperm, but in the Comstockiella type some paternal chromosomes are lost from the soma and under the diaspidid type, the full paternal haplotype is discarded early in embryonic development (Brown 1965). In other words, in the earliest form of paternal genome elimination, all paternal chromosomes are maintained in the soma but in later evolving types, some or all paternal chromosomes are eliminated early on in development. Early molecular work suggested that there have been multiple independent transitions in type of paternal genome elimination within armored scales (Morse and Normark 2006), but we lack the taxon sampling of this group to confirm or refute this specific result. We are, however, able to track the fate of the X chromosome across this transition.

In sharp contrast to the loss of the X in Monophlebidae, the X chromosome is almost entirely intact in the early diverging paternal genome elimination species, starting with the crapemyrtle bark scale, *Acanthococcus lagerstroemiae*. Indeed, while there have been some autosome-autosome rearrangements between these two species, the overall karyotype has not changed drastically (n = 10 vs. n = 9) and at least four chromosomes, including the former-X, are largely intact across the transition. Within the paternal genome elimination clade, complex rearrangements appear even less common. The former-X in particular does not undergo any major changes until the soft scale *Toumeyella liriodendri*, in which it splits into two chromosomes along with most other chromosomes as part of a large increase in chromosome number. Only in the family Pseudococcidae, the true mealybugs, has there been substantial rearrangement to the chromosomes, including former-X-linked orthologs, which are scattered across all five autosomes.

In keeping with the modest chromosomal changes, we recovered far fewer differences at the level of individual gene conservation or novelty. Indeed, there were only two reliably X-linked orthologs lost in the transition to paternal genome elimination, and fewer than 100 orthogroups gained. The two lost orthologs were a PITH domain-containing protein and CCN family member 1. Intriguingly, the latter was also identified as lost from the hermaphrodite group. This pattern could hint at a shared molecular mechanism across the two losses or be mere coincidence. As discussed above,there is currently no evidence for orthologs of this protein playing a sex determining role in other insects, though there is little work on invertebrates. The ortholog ID came from a BLAST hit to the whitefly *Bemesia tabaci*, a haplodiploid hemipteran pest (Blackman and Cahill 1998). A targeted BLAST of the *A. pisum* annotation recovered two similarly annotated orthologs, but both are located on autosomes. In other words, an ortholog of this protein was conserved across another loss of X sex determination and is not X-linked in other Sternorrhyncha, so its status as a sex determination gene in scale insects is dubious. Still, as a singular target for further study, it is quite tractable.

Turning to orthogroups gained in the transition, many of these have no currently defined function, so their significance to sex determination and reproduction remains to be seen. In contrast to the orthogroups gained in the hermaphrodite lineage, far fewer had no BLAST hits at all. Part of this can be attributed to the number of species compared, but it is also worth noting that mealybugs are by far the best-studied scale insects, so there is more genetic data for them (de la Filia et al. 2021; Liesenfelt et al. 2025b; Mongue et al. 2025; Vea et al. 2025). While most genes have yet to be characterized, these putatively unique genes make an intriguing set of targets for future study with the rise of functional genomics techniques like RNAi in mealybugs (Khan et al. 2018; Bansal and Fernandez 2025). For now, we can conclude that like in Monophlebidae, sex determination turnover is not associated with any appreciable gene loss. This pattern is also visible in the distribution of sex-biased genes across the genome, as discussed below.

### State of the X before turnover and the drivers of its change

A study on other independently evolved paternal genome elimination systems (Herbette and Ross 2023) has suggested that transitions from X sex determination may be easier if a larger portion of the genome is X-linked (Haig 1993). Previous research has shown a lower chromosome count is associated with the evolution of paternal genome elimination in mites (Blackmon et al. 2015) and more recent research has shown an association between a larger X and paternal genome elimination in fungus gnats and springtails, in which males are somatically diploid with the exception of the X and paternal chromosomes are only eliminated in the germline (Anderson et al. 2022). At first glance, this scenario seems plausible with the hermaphroditic scales; the number of chromosomes decreases in closer relatives to the hermaphrodites. However, the reduction in karyotype is not associated with an increase in X-linked genes, as *C. castanopsis* has a roughly comparable, but slightly lower proportion of X-linked genes than *N. texana*. The pattern is even less convincing for the paternal genome elimination scale insects, which did not see a substantial chromosome number change across the transition, nor is the former-X a particularly large or gene-rich chromosome. So what distinguishes turnover in scale insects from other systems?

To better understand the dynamics of sex determination turnover, we further characterized the X chromosome of *N. texana.* First, we looked for evidence of how the X chromosome is regulated in scale insects. Briefly, because the autosomal genome is diploid in both sexes but the X is diploid in females and haploid in males, gene expression levels may need to be compensated in males (Disteche 2012; Gu and Walters 2017). We found that the haploid X in males expresses genes at the same level as the diploid autosomes; in other words, the dosage is compensated, as seen in other X sex chromosome Hemiptera (Jaquiéry et al. 2013; Pal and Vicoso 2015). This simple observation has two implications. To start, under haplodiploidy, the autosomes and former-X have the same ploidy, so the gene dosage is the same across the genome. Second, thanks to this new parity, the former-X may have evolved to be down-regulated to match the expression levels of the now single-copy autosomes in males. If so though, it would require changes in regulation only on the former-X.

Next, we explored the sex-biased expression profiles of genes on the X and autosomes. Sex chromosome theory offers two distinct predictions for such genes. In general, because of the ploidy difference between the sexes, the X chromosome spends more time over the generations in females than in males, and is predicted to accumulate female-biased genes (Klein et al. 2021). However, because genes on the X expressed in males are expressed in a haploid state, they are exposed to more efficient selection (Charlesworth et al. 1987), so male-biased genes may accumulate on the X, especially if their fitness effects are recessive (Rice 1984). In practice, the former outcome seems more common, as X chromosomes are often feminized (Meisel et al. 2012; Allen et al. 2013; Albritton et al. 2014), and only rarely observed to be masculinized (Jaquiéry et al. 2013; Robert B. Baird et al. 2025). Our observations from *N. texana* fall into the latter set: a masculinized X chromosome relative to autosomes. It is worth noting that both of the other examples occur in systems with unusual transmission of the chromosomes (Jaquiéry et al. 2013; Baird et al. 2023), which differ from the presumed Mendelian inheritance patterns of *N. texana*.

It may simply be that male-benefitting alleles are, on average, more recessive than female-benefitting alleles in this species, as Rice (1984) would predict, but there may be other contributing factors as well. Chiefly, sex-biased gene expression is extreme across the entire genome, with even the autosomes of *N. texana* having a majority of female-biased genes based on our analyses. This pattern itself is not unprecedented, as other insects with extreme morphological sexual dimorphism also have highly dimorphic patterns of gene expression (Mongue et al. 2025; Mongue et al. 2025), but it may complicate selective dynamics. If sexual dimorphism is high and more genes are female-biased in expression (a proxy for fitness benefit), then many male-biased alleles will be in close proximity to sexually-antagonistic loci by chance. This could hinder the fixation of genes with sex-limited or sexually-antagonistic fitness effects if they end up locked in inversions together (McAllester and Pool 2025) or even merely in close proximity and in linkage disequilibrium (Patten et al. 2010). This potential problem should be present across all chromosomes, but the X sex chromosome could offer some reprieve.

Because of its haploid expression in males, the X should offer more efficient selection on male-biased alleles than any other part of the genome (Charlesworth et al. 1987). Under these conditions, the X may be the best place in the genome for selection on male-biased traits. Additional gene expression analyses will be required to test this prediction across scale insects and other taxa, as this logic should hold for any highly sexually dimorphic system with sex chromosomes.

Regardless of the cause of the gene expression patterns above, they help explain the loss of X sex determination. If sex determination system turnover is driven by sexual conflict between chromosomes (van Doorn and Kirkpatrick 2007) and haplodiploidy represents a state in which the whole genome is biased towards females (Blackmon et al. 2015; Klein et al. 2021; Hitchcock et al. 2022), then it stands to reason that its evolution would be more favored in systems with a preponderance of female-biased genes distributed throughout the genome. In other words, it may be that sex determination turnover in scale insects was driven, or at least enabled, by extreme sexual dimorphism, which is a common feature across this superfamily (Vea et al. 2016). Once this transition occurred, however, there would be no further pressure for a preferential accumulation of sex-biased alleles in one part of the genome in particular, as every chromosome has the same transmission dynamics. Fitting with this prediction, the biased distribution of genes does appear to persist on the former-X of *E. pela*, with a continued excess of male-biased genes and relative dearth of female-biased genes. Only later, in *T. liriodendri*, does this pattern break down as the former-Xs undergo more rearrangement and the composition of sex-biased genes evens out. Future expression analyses across other X chromosome scale insects and paternal genome elimination species will be necessary to confirm the robustness of this pattern.

### Other potential drivers of change

One final consideration is that scale insects, like other phloem-feeding Hemiptera, employ endosymbiotic organisms, often bacteria, to provision essential nutrients (Baumann 2005). These symbionts are transmitted from mothers to offspring but are purged from males before adulthood (Kono et al. 2008). In other words, males are an evolutionary dead end for their symbionts. Because of this transmission dynamic, symbionts could increase their evolutionary fitness by skewing sex ratios towards females (Ross et al. 2012b). Indeed, consider that hermaphroditic monophlebids are phenotypically female (with the addition of testis tissue, Johnston 1912) and pass on endosymbiotic bacteria to their hermaphroditic offspring (Szklarzewicz et al. 2020). Though the exact developmental sequence remains unclear, the sperm-producing portion of the ovotestis gonad of hermaphrodites appears to arise from supernumerary sperm nuclei that persist in the embryonic cytoplasm (Royer 1975). Royer (1975) also observed that the endosymbiotic bacteria physically associate with these sperm nuclei, as if supporting them. Thus, the endosymbiotic bacteria appear to have both plausible selective pressure and opportunity to capture sex determination in Monophlebidae. In fact, this hypothesis was first suggested some 20 years ago (Normark 2004). Similar arguments apply to paternal genome elimination species, in which sex determination of embryos appears to be labile, even across an individual’s lifespan (Ross et al. 2011). We leave deeper exploration of symbionts for a future study, but our new sequencing data offers new chances to characterize symbionts in previously unstudied scale insects.

## Conclusions

We generated 17 assemblies for previously unsequenced scale insects to explore the change in sex determination across this group. We found two transitions away from X chromosome sex determination, once for simultaneous hermaphroditism and once for paternal genome elimination haplodiploidy. These two transitions took very different forms at the genomic level, with the former associated with a reduction in karyotype and extensive rearrangements and the latter transition involving minimal change to the X chromosome at first. We found few to no individual X-linked genes lost in these transitions, suggesting that the genetic machinery for sex determination was supplanted rather than lost outright. The one exception, an ortholog of CCN family 1 proteins, does not appear crucial to sex determination in other Hemiptera but makes a suitable target for future study. Finally, in characterizing the X chromosome before the transition, we found a surprisingly masculinized chromosome compared to an overall feminized genome. We suggest that extreme sexual dimorphism and widespread sexual conflict across the genome may be a driving force of sex determination turnover in scale insects. We hope these new results and data are useful for the study of scale insects in both basic and applied settings, and more generally, that they encourage other studies of sex determination turnovers across non-model systems.

## Acknowledgements

We are grateful to Douglass Miller and Zachary Lahey for their constructive comments on earlier drafts of the manuscript. We thank the following contributors for supplying invaluable material: Douglass Miller and Barbara Denno for *Crypticerya rileyi, Orthezia annae*, *Puto cupressi*, *Puto decorosus*, and *Stomacoccus platani*, Mark Zenoble for *Crypticerya genistae*, Elena Oey for the *Puto* sp., Edward Ruden for *Pityococcus rugulosus*, Xinyi Zheng for *Neogreenia osmanthus*, and Lyle Buss for *Eumargarodes laingi*. The Florida Department of Agriculture and Consumer Services, Division of Plant Industry (FDACS-DPI) supported this work. These analyses were enabled by the infrastructure of the University of Florida Information Technology Research Computing (UFIT-RC)’s HiPerGator computing cluster.

