## Supplemental figures and tables for "Leaving X: How scale insects evolved alternatives to chromosomal sex determination"

**Leaving X Supplementary information**

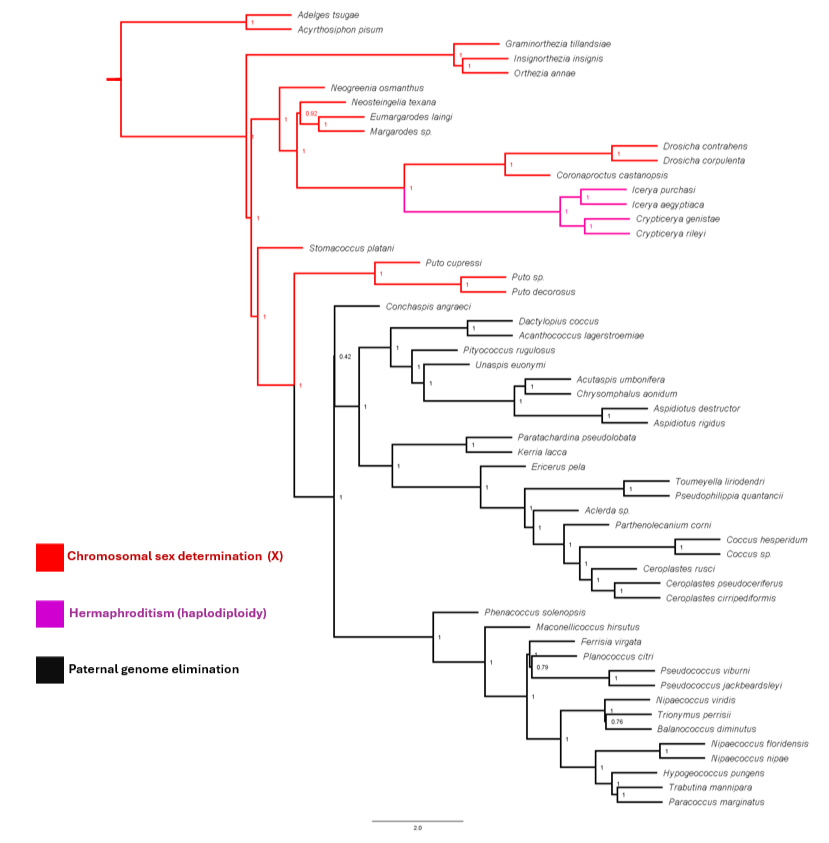

**Figure S1.** Consensus species tree from gene trees recapitulates relationships from ML tree. Node values indicate posterior quartet support values and show high confidence across the shifts in sex determination. As in the main text, branches follow the sex determination color scheme: red for X sex chromosomes, purple for hermaphrodites, and black for PGE.

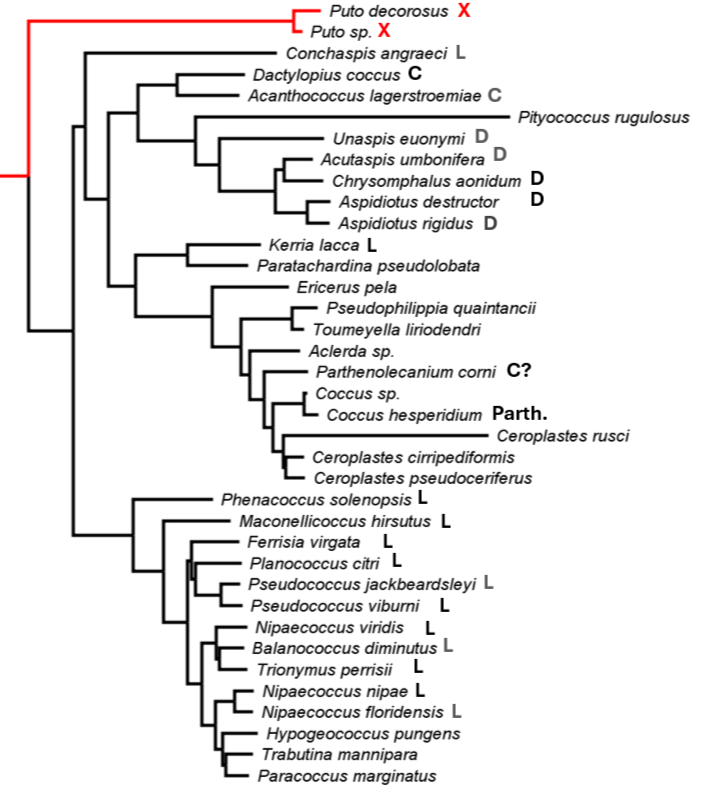

**Figure S2.** Mapping types of paternal genome elimination to the scale insect phylogeny. Paternal genome elimination types are abbreviated as L for lecanoid, C for Comstockiella, or D for diaspidid. Parth. is an abbreviation of parthenogenesis. Labels in full black represent species-specific reports of chromosome behavior, those in grey have data from other species in the same genus, and those without a label have no available data. Data come from records collected in Gavrilov, 2007.

| Species | Family | Sex Determination System | Karyotype (n) | Assembly Accession | Assembly Reference | Sex determination reference |
| --- | --- | --- | --- | --- | --- | --- |
| *Planococcus citri* | Pseudococcidae | PGE-L (C) | 5^a^ | GCA_950023065.1 | (Ross et al. 2024) |  |
| *Nipaecoccus viridis* | Pseudococcidae | PGE-L | 5^a^ | GCA_052327235.1 | (Liesenfelt et al. 2025) |  |
| *Toumeyella liriodendri* | Coccidae | PGE | 17^a^ | GCA_041937245.1 | (Mongue et al. 2024a) |  |
| *Icerya purchasi* | Monophlebidae | Androdioecy (haplodiploidy) | 2^a^ | GCA_952773005.1 | (Mongue et al. 2024b) | (Mongue et al. 2021) |
| *Coronaproctus castanopsis* | Monophlebidae | XX/X0(I*) | 3^a^ | GCA_032883995.1 | (Huang et al. 2024) | This study |
| *Acyrthosiphon pisum* | Aphididae (outgroup) | XX/X0 | 4^a^ | GCF_005508785.2 |  |  |
| *Adelges tsugae* | Adelgidae (outgroup) | X1X1X2X2/X1X2 | 10^a^ | GCA_045999835.2 |  |  |
| *Margarodes* sp. | Margarodidae | X sex chromosomal | ? | [SRA] |  |  |
| *Drosicha corpulenta* | Monophlebidae | X sex chromosomal | ? | SRA |  |  |
| *Icerya aegyptiaca* | Monophlebidae | Androdioecy (haplodiploidy) | 2 | Different paper SRA |  |  |
| *Phenacoccus solenopsis* | Pseudococcidae | PGE-L | 5^a^ | GCA_009761765.1 | (Li et al. 2020) | (Li et al. 2020) |
| *Maconellicoccus hirsutus* | Pseudococcidae | PGE-L | 5 | GCA_900064465.1 |  | (Nur et al. 1987) |
| *Ferrisia virgata* | Pseudococcidae | PGE-L | 5 | GCA_900060175.1 |  | (Nur et al. 1987) |
| *Pseudococcus viburni* | Pseudococcidae | PGE-L | 5 | PRJEB47083 | (Vea et al. 2025) | (Schrader 1923) |
| *Pseudococcus jackbeardsleyi* | Pseudococcidae | PGE (I) | 5^a^ | GCA_038380155.1 |  |  |
| *Balanococcus diminutus* | Pseudococcidae | PGE (I) | 5^a^ | PRJEB64510 |  |  |
| *Trionymus perrisii* | Pseudococcidae | PGE-L | 8 | GCA_900050545.1 |  | (Gavrilov 2004) |
| *Hypogeococcus pungens* | Pseudococcidae | PGE (I) |  | GCA_018107765.1 |  |  |
| *Trabutina mannipara* | Pseudococcidae | PGE (I) |  | GCA_900080175.1 |  |  |
| *Paracoccus marginatus* | Pseudococcidae | PGE (I) |  | GCA_900065295.1 |  |  |
| *Dactylopius confusus* | Dactylopiidae | PGE-C | 5 | SRR1821914 |  | (Aquino et al. 1994) |
| *Acanthococcus lagerstroemiae* | Eriococcidae | PGE (I) | 9^a^ | GCA_031841125.1 |  | (Gavrilov 2007) |
| *Unaspsis euonymi* | Diaspididae | PGE-D | 4 | SRR1821978 |  | (Brown 1965) |
| *Aspidiotus destructor* | Diaspididae | PGE-D | 4 | GCA_035079395.1 |  | (Brown 1965) |
| *Aspidiotus rigidus* | Diaspididae | PGE-D (I) |  | GCA_035079385.1 |  | (Gavrilov 2007) |
| *Chrysomphalus aonidum* | Diaspididae | PGE-D | 4 | SRR1821907 |  | (Brown 1965) |
| *Acutaspis umbonifera* | Diaspididae | PGE-D (I) |  | SRR1821893 |  | (Gavrilov 2007) |
| *Kerria lacca* | Kerriidae | PGE-L | 9 | SRS525030 |  | (TSS 1964) |
| *Paratachardina pseudolobata* | Kerriidae | PGE (I) |  | SRR5320111 |  |  |
| *Ericerus pela* | Coccidae | PGE | 9^a^ | GCA_011428145.1 |  |  |
| *Aclerda* sp. | Aclerdidae | PGE |  | SRR1821892 |  | This study |
| *Parthenolecanium corni* | Coccidae | PGE-C |  | GCA_038050395.1 |  | (Gavrilov and Kuznetsova 2004) |
| *Coccus* sp. | Coccidae | PGE (I) |  | SRR1821911 |  |  |
| *Coccus hesperidum* | Coccidae | Parthenogenesis (Deuterotoky) | 7^a^ | GCA_964257065.1 |  |  |
| *Ceroplastes rusci* | Coccidae | PGE (I) |  | GCA_050859255.1 |  |  |
| *Ceroplastes pseudoceriferus* | Coccidae | PGE (I) | 18^a^ | GCA_050872025.1 |  |  |
| *Ceroplastes cirripediformis* | Coccidae | PGE (I) |  | SRR1821905 |  |  |
| *Drosicha contrahens* | Monophlebidae | XX/X0 | 3^a^ | GCA_054252755.1 |  |  |

**Table S1. List of species and sex determination systems for taxa downloaded from public repositories.** For the listed sex determination system, paternal genome elimination is abbreviated PGE, with an additional qualifier of L (lecanoid), C (Comstockiella), or D (diaspidid) whenever information is available. Each species is additionally qualified with either “C” for confirmed for published karyotypes or direct analysis of sequence data in this study, or “I” for inferred when sex determination system was only available for other members of the genus or family. We note the special case of *Coronaproctus castanopsis*, which we infer to be XX/X0 based on several lines of evidence, with an *. For Karyotype information, where information is available, species with chromosome level references are denoted an an ^a^, otherwise information comes from published karyotype.

| Species | Family | Sex Determination System | Karyotype (n) | Accession | Sex determination reference |
| --- | --- | --- | --- | --- | --- |
| *Graminorthezia tillandsiae* | Ortheziidae | X sex chromosomal (C) | 7^a^ | [to be added] | This study |
| *Orthezia annae* | Ortheziidae | X sex chromosomal (I) | ? |  | (Gavrilov 2004) |
| *Insignorthezia insignis* | Ortheziidae | X sex chromosomal (I) | ? |  | (Gavrilov 2007) |
| *Neogreenia osmanthus* | Qinococcidae | ?? (Inferred X) | ? |  | This study |
| *Eumargarodes laingi* | Margarodidae | X sex chromosomal (I) | ? |  | (Gavrilov 2007) |
| *Neosteingelia texana* | Kuwaniidae | X sex chromosomal (C) | 9^a^ |  | This study |
| *Stomacoccus platani* | Steingeliidae | X sex chromosomal (C) | ? |  | This study |
| *Puto decorosus* | Putoidae | X sex chromosomal (C) | ? |  | This study |
| *Puto cupressi* | Putoidae | X sex chromosomal (C) | ? |  | This study |
| *Puto* sp. | Putoidae | X sex chromosomal (C) | 10^a^ |  | This study |
| *Nipaecoccus nipae* | Pseudococcidae | PGE (C) | 5 |  | (Nur et al. 1987) |
| *Nipaecoccus floridensis* | Pseudococcidae | PGE (I) | ? |  | (Gavrilov 2007) |
| *Conchaspis angraeci* | Conchaspidae | PGE (I) | ? |  | (Brown 1959) |
| *Pseudophilippia quantancii* | Coccidae | PGE (I) | ? |  | (Gavrilov 2007) |
| *Pityococcus rugulosus* | Pityococcidae | ?? (Inferred PGE) | ? |  | This study |

**Table S2. List of species and sex determination systems for taxa sequenced *de novo* for this study.** For the listed sex determination system, each is qualified with either “C” for confirmed for published karyotypes or direct analysis of sequence data in this study, or “I” for inferred when sex determination system was only available for other members of the genus or family. Thus by this heuristic, *Neogreenia osmanthus* has no supporting information, as the sole member of a newly erected family. We tentatively assign it as X chromosomal based on its phylogenetic placement with other X chromosomal systems. For karyotype information, where information is available, species with chromosome level references are denoted an an ^a^, otherwise information comes from published karyotype.

|  | *Graminorthezia tillandsiae* | | | | *Neosteingelia texana* | | | | *Crypticerya genistae* | | | | *Crypticerya rileyi* | | | | *Puto sp.* | | | |
| --- | --- | --- | --- | --- | --- | --- | --- | --- | --- | --- | --- | --- | --- | --- | --- | --- | --- | --- | --- | --- |
| Assembly Statistics | Primary HiFi Assembly (Hifiasm) | Purged HiFi Assembly (Purge_dups, two rounds) | HiC Scaffolded Assembly (YaHs) | Curated Assembly (Juicebox) | Primary HiFi Assembly (Hifiasm) | Purged HiFi Assembly (Purge_dups) | HiC Scaffolded Assembly (YaHs) | Curated Assembly (Juicebox) | Primary HiFi Assembly (Hifiasm) | Purged HiFi Assembly (Purge_dups) | HiC Scaffolded Assembly (YaHs) | Curated Assembly (Juicebox) | Primary HiFi Assembly (Hifiasm) | Purged HiFi Assembly (Purge_dups) | HiC Scaffolded Assembly (YaHs) | Curated Assembly (Juicebox) | Primary HiFi Assembly (Hifiasm) | Purged HiFi Assembly (Purge_dups) | HiC Scaffolded Assembly (YaHs) | Curated Assembly (Juicebox) |
| Total length (bp) | 443,629,712 | 166,095,113 | 166,172,613 | 166,173,213 | 1,022,380,005 | 947,543,051 | 947,592,751 | 947,593,051 | 1,204,795,380 | NA | 1,204,837,080 | (No further curation needed) | 1,876,127,373 | 1,719,634,221 | 1,876,216,173 | 1286973622 | 345,955,981 | 326,410,667 | 326,579,867 | 326,583,867 |
| Sequence count (n) | 4,484 | 1,094 | 319 | 313 | 1,416 | 629 | 388 | 385 | 1,224 | NA | 934 |  | 9,590 | 7,590 | 9,180 | 2 | 2,813 | 2,168 | 672 | 638 |
| N50 (bp) | 175,098 | 256,000 | 19,678,824 | 19,678,824 | 3,613,973 | 3,772,205 | 94,726,905 | 95,070,468 | 6,803,401 | NA | 649,014,850 |  | 2,094,200 | 2,346,537 | 576,146,302 | 710,827,320 | 276,895 | 293,500 | 29,211,445 | 30,115,152 |
| L50 (n) | 778 | 217 | 3 | 3 | 91 | 81 | 4 | 4 | 55 | NA | 0 |  | 235 | 201 | 1 | 0 | 346 | 313 | 4 | 4 |
| GC content (%) | 33.5 | 33.6 | 33.6 | 33.6 | 32.1 | 32.2 | 32.2 | 32.2 | 33.2 | NA | 33.2 |  | 33.5 | 33.3 | 33.5 | 31.9 | 32.3 | 32.2 | 32.2 | 32.2 |
| N content (%) | 0 | 0 | 0.047 | 0.047 | 0 | 0 | 0.005 | 0.005 | 0 | NA | 0.003 |  | 0 | 0 | 0.005 | 0.005 | 0 | 0 | 0.053 | 0.053 |
| Assembly BUSCO completeness  (hemipteraodb10, n=2510) | C:89.8%[S:6.5%,D:83.3%],F:2.0%,M:8.2% | C:88.6%[S:82.6%,D:6.0%],F:2.2%,M:9.2% | C:89.5%[S:83.9%,D:5.7%],F:0.9%,M:9.6% | C:89.5%[S:83.9%,D:5.7%],F:0.9%,M:9.6% | C:88.4%[S:83.2%,D:5.2%],F:1.2%,M:10.4% | C:88.5%[S:85.9%,D:2.5%],F:1.3%,M:10.2% | C:88.7%[S:86.2%,D:2.5%],F:1.2%,M:10.1% | C:88.6%[S:86.2%,D:2.5%],F:1.2%,M:10.2% | C:88.2%[S:84.7%,D:3.5%],F:2.5%,M:9.3% | NA | C:88.0%[S:85.6%,D:2.4%],F:1.2%,M:10.7% |  | C:96.1%[S:23.1%,D:73.0%],F:1.4%,M:2.5% | C:96.1%[S:29.8%,D:66.3%],F:1.4%,M:2.5% | C:96.1%[S:28.0%,D:68.0%],F:0.5%,M:3.4% | C:87.0%[S:85.3%,D:1.7%],F:1.1%,M:11.9% | C:87.9%[S:82.8%,D:5.1%],F:1.5%,M:10.6% | C:87.8%[S:85.5%,D:2.3%],F:1.4%,M:10.7% | C:88.4%[S:86.1%,D:2.2%],F:1.2%,M:10.4% | C:88.4%[S:86.2%,D:2.2%],F:1.2%,M:10.4% |
| Gene annotation (BRAKER protein) | NA | NA | NA | 14,642 | NA | NA | NA | 18,697 | NA | NA | 20,330 |  |  |  |  | 19,871 |  |  |  | 16,042 |
| Annotation BUSCO completeness | NA | NA | NA | C:92.4%[S:85.0%,D:7.4%],F:0.6%,M:6.9% | NA | NA | NA | C:86.5%[S:82.8%,D:3.7%],F:3.0%,M:10.4% | NA | NA | C:44.9%[S:43.0%,D:1.8%],F:4.2%,M:50.9% |  |  |  |  | C:84.5%[S:81.5%,D:3.1%],F:2.7%,M:12.7% |  |  |  | C:91.3%[S:87.7%,D:3.6%],F:1.4%,M:7.3% |

**Table S3. Assembly statistics of five new chromosome-level scale insect assemblies.**

| Assembly Statistics | *Insignorthezia insignis* | *Orthezia annae* | *Neogreenia osmanthus* | *Eumargarodes laingi* | *Puto decorosus* | *Conchaspis agraeci* | *Pityococcus rugulosus* | *Pseudophilippia quantancii* | *Nipaecoccus nipae* | *Nipaecoccus floridensis* | *Stomacoccus platani* |
| --- | --- | --- | --- | --- | --- | --- | --- | --- | --- | --- | --- |
| Assembler | Megahit | Megahit | Megahit | Megahit | Megahit | Megahit | SPAdes | Megahit | Megahit | Megahit | Hifiasm + purge_dups |
| Total length (bp) | 132,904,464 | 98,825,374 | 437,436,217 | 468,506,254 | 323,404,211 | 253,487,361 | 502,548,312 | 479,091,804 | 203,613,691 | 242,050,423 | 84,070,446 |
| Sequence count (n) | 19,532 | 25,664 | 445,232 | 393,978 | 120,914 | 36,577 | 725,665 | 298,511 | 64,387 | 71,668 | 1,252 |
| N50 (bp) | 27,356 | 17,330 | 1,487 | 1,729 | 7,295 | 32,604 | 1,808 | 2,628 | 10,871 | 7,454 | 101,332 |
| L50 (n) | 1,379 | 1,560 | 79,517 | 71,798 | 11,359 | 1,951 | 62,286 | 50,786 | 4,512 | 8,908 | 249 |
| GC content (%) | 31.7 | 29.2 | 33.9 | 39.1 | 32.2 | 26.2 | 37.1 | 35.6 | 35.7 | 35.1 | 38.1 |
| Assembly BUSCO completeness  (hemipteraodb10, n=2510) | C:87.5%[S:86.1%,D:1.4%],F:3.4%,M:9.1% | C:87.7%[S:86.0%,D:1.7%],F:3.5%,M:8.8% | C:49.9%[S:47.8%,D:2.1%],F:18.7%,M:31.4% | C:41.6%[S:28.1%,D:13.5%],F:19.6%,M:38.8% | C:66.3%[S:64.5%,D:1.8%],F:11.7%,M:22.0% | C:85.3%[S:83.3%,D:2.0%],F:4.0%,M:10.7% | C:57.5%[S:50.6%,D:6.9%],F:21.2%,M:21.4% | C:58.2%[S:55.1%,D:3.1%],F:14.3%,M:27.5% | C:87.0%[S:84.7%,D:2.3%],F:5.2%,M:7.8% | C:84.9%[S:82.5%,D:2.4%],F:5.6%,M:9.5% | C:82.4%[S:72.5%,D:9.9%],F:1.4%,M:16.3% |

**Table S4. Assembly statistics of 11 new draft scale insect assemblies.** Species were sequenced as 150bp paired-end Illumina reads and assembled with Megahit. Due to low contiguity, these assemblies were not archived, only their raw reads, as listed in table S2 above.

*Identification of sex chromosomes*

Across the seven chromosomal scaffolds of *G. tillandsiae*, male read coverage ranged from 52.8x to 107.3x. Across six of the seven, that range narrowed to 96.6x - 107.3x, thus that single scaffold (scaffold_2) with 52.8x coverage is clearly the X chromosome at half coverage in males. Incidentally, the X is the largest chromosomal scaffold in the assembly (at 26.6Mb).

For *N. texana*, male read coverage was lowest on scaffold_1 at 21.5x. Across the other 8 chromosomal scaffolds, coverage ranged from 32.6x - 33.8x. Thus, although one single scaffold had clearly female-biased coverage, it was marginally higher than the naive expectation. Further investigation of coverage along the length of the scaffold suggested that the ends of the scaffold had higher coverage (~25x) than the bulk of the middle which showed the expected ~16x coverage. [SHOW IT]. We did not observe any obvious misassemblies in the Juicebox curation, and the higher coverage does not reach the full diploid expectation either. Overall, coverage was consistent with an X chromosomal system, but the coverage variation on the X is suggestive of either an excess of repetitive sequences on the ends of the X or, potentially, a Y chromosome with some lingering homology to portions of the X. Direct karyotyping will be required to confirm or refute the latter.

*Annotation*

In both species we tested the utility of the new *ab initio* annotation software helixer (Holst et al. 2023) against more traditional methods using BRAKER with protein evidence (Hoff et al. 2019). Starting with *G. tillandsia*, we annotated 12,211 protein coding genes with helixer, which included 83.8% of expected hemipteran BUSCOs. We annotated more genes, 14,642, with a higher BRAKER, score of 92.4%. Turning to *N. texana*, with helixer, we identified 18,679 protein coding genes that covered 69.1% of expected BUSCOs. BRAKER recovered a similar 18,697 genes, but with a BUSCO score of 86.5%. In both cases, we selected the BRAKER annotations for downstream analyses. It is also worth noting that as a deep learning tool, helixer’s algorithm is not tunable beyond the initial starting model weights (Holst et al. 2023). While perhaps not surprising that helixer performed worse than BRAKER with protein evidence. What is more surprising is that for *N. texana*, the number of annotated genes differed by only 12, yet the BUSCO scores were markedly worse, suggesting that the helixer annotations were less accurate overall.

| Gene  (*G. tillandsiae*) | Top BLAST hit |
| --- | --- |
| Gt_scaffold_2-_g4535.t1 | tumor protein p63-regulated gene 1 protein-like isoform X2 [Planococcus citri] |
| [Gt_scaffold_2+_g3869.t1](https://blast.ncbi.nlm.nih.gov/Blast.cgi#) | leucine-rich repeat neuronal protein 1-like isoform X2 [Planococcus citri] |
| Gt_scaffold_2+_g3779.t1 | rhythmically expressed gene 5 protein [Adelges cooleyi] |
| Gt_scaffold_2-_g4539.t1 | C1 family peptidase 26-29-p isoform X2 [Halictus rubicundus] |
| Gt_scaffold_2-_g3776.t2 | arf-GAP with Rho-GAP domain, ANK repeat and PH domain-containing protein 1 [Planococcus citri] |
| Gt_scaffold_2+_g3879.t1 | No hits |
| Gt_scaffold_2+_g4377.t1 | basic proline-rich protein-like [Macrosteles quadrilineatus] |
| **Gt_scaffold_2+_g3608.t1** | **CCN family member 1 [Bemisia tabaci]** |

**Table S5. X-linked orthologs missing in hermaphrodites: *I. purchasi, C. genistae,* and *C. rileyi*.**

| Gene  (*G. tillandsiae*) | Top BLAST hit |
| --- | --- |
| [**Gt_scaffold_2+_g3608.t1**](https://blast.ncbi.nlm.nih.gov/Blast.cgi#) | **CCN family member 1 [Bemisia tabaci]** |
| Gt_scaffold_2-_g3491.t1 | PITH domain-containing protein GA19395 isoform X2 [Lycorma delicatula] |

**Table S6. X-linked orthologs missing in paternal genome elimination species.**

**Table S6. Excel file of BLAST hits comparing X species and hermaphrodites.**

**Table S7. Excel file of BLAST hits comparing X species and paternal genome elimination species.**
